# Telomeric small RNAs in the genus *Caenorhabditis*

**DOI:** 10.1101/612820

**Authors:** Stephen Frenk, Evan H. Lister-Shimauchi, Shawn Ahmed

## Abstract

Telomeric DNA is composed of simple tandem repeat sequences and has a G-rich strand that runs 5’ to 3’ towards the chromosome terminus. Small RNAs with homology to telomeres have been observed in several organisms and could originate from telomeres or from interstitial telomere sequences (ITSs), which are composites of degenerate and perfect telomere repeat sequences found on chromosome arms. We identified *C. elegans* small RNAs composed of the *Caenorhabditis* telomere sequence (TTAGGC)_n_ with up to three mismatches, which might interact with telomeres. We rigorously defined ITSs for genomes of *C. elegans* and for two closely related nematodes, *C. briggsae* and *C. remanei*. We found that most telomeric small RNAs with mismatches originated from ITSs, which were depleted from mRNAs and but were enriched in introns whose genes often displayed hallmarks of genomic silencing. *C. elegans* small RNAs composed of perfect telomere repeats were very rare but were increased by several orders of magnitude in *C. briggsae* and *C. remanei*. Major small RNA species in *C. elegans* begin with a 5’ guanine nucleotide, which was strongly depleted from perfect telomeric small RNAs of all three *Caenorhabditis* species. Perfect telomeric small RNAs corresponding to the G-rich strand of the telomere commonly began with 5’ UAGGCU and 5’UUAGGC, whereas C-rich strand RNAs commonly begin with 5’CUAAGC. In contrast, telomeric small RNAs with mismatches had a mixture of all four 5’ nucleotides. Together, our results imply that perfect telomeric small RNAs have a mechanism of biogenesis that is distinct from known classes of small RNAs and that a dramatic change in their regulation occurred during recent *Caenorhabditis* evolution.

## Introduction

Telomeres are repetitive sequences at the end of linear eukaryotic chromosomes that possess a G-rich strand that runs 5’ to 3’ towards chromosome termini (Smogorzewska and de Lange, 2004). These sequences play an essential role in the maintenance of genome stability by protecting chromosomes ends from recognition by DNA damage response pathways, thereby preventing chromosome fusions (Palm and de Lange, 2008). Due to the end replication problem, telomeres are eroded with each cell division. Persistent telomere erosion triggers DNA damage response activation and results in senescence. To counteract telomere erosion, the reverse transcriptase telomerase uses an RNA template to add *de novo* telomeric repeats to chromosome termini. Although telomerase is essential for maintenance of proper germline function across generations, this enzyme is absent from most human somatic cells. Somatic silencing of telomerase and associated telomere shortening represents a barrier to tumorigenesis in large mammals (Gomes et al., 2011; Seluanov et al., 2007). Telomere maintenance in around 90% of cancers is achieved through activation of telomerase, while the remaining 10% utilize a telomerase-independent mechanism that is less well understood known as Alternative Lengthening of Telomeres (ALT) (Pickett and Reddel, 2015).

The protective function of telomeres depends in part on binding of a protein complex known as Shelterin, composed of POT1, TRF1, TRF2, TIN2, TPP1 and RAP1, that interacts with single-(POT1) and double-stranded (TRF1 and TRF2) telomeric DNA to promote telomere capping as well as the maintenance of a repressive chromatin environment (de Lange, 2018; Dyer et al., 2017). Heterochromatin associated histone modifications such as H3K9me3, H3k20me3 and hypoacetlyation of H3 and H4 are a common feature of telomeres (Schoeftner and Blasco, 2010). Somewhat paradoxically, transcription by RNA polymerase II has also been observed at telomeres, leading to formation of the G-rich telomere repeat containing long non-coding RNA TERRA, which is composed of subtelomeric DNA sequences and some perfect telomere repeats, as well as the less abundant C-rich antisense telomeric transcript ARRET (Azzalin et al., 2007; Bah et al., 2012). TERRA expression is upregulated at critically short and damaged telomeres, where it may recruit telomerase and chromatin modifiers in order to promote telomere elongation and protect telomeres (Azzalin and Lingner, 2015; Cusanelli et al., 2013). It has been observed that TERRA prominently coats telomeres of the silent X chromosome in mammals and may promote X chromosome inactivation, but also localizes to autosomal telomeres and to non-telomeric regions of the genome (Chu et al., 2017; Schoeftner and Blasco, 2008; Zhang et al., 2009).

Silencing of certain repetitive regions of the genome such as pericentromeres have been shown to require transcription of long non-coding transcripts that interact with homologous small interfering RNAs (siRNAs) that are bound to Argonaute proteins (Holoch and Moazed, 2015; Reinhart and Bartel, 2002). siRNAs can also promote mRNA destruction or translational repression in the cytoplasm.

Endogenous small RNA pathways have been well characterized in the nemotode worm *Caenorhabditis elegans* (*C. elegans*) (Billi et al., 2014). Primary siRNAs can be created by DCR-1, the *C. elegans* homologue of the highly conserved RNAse class III enzyme Dicer, which cleaves double stranded RNA transcripts to create 26 nucleotide long siRNAs beginning with a 5’ guanine nucleotide (26G RNAs) (Bernstein et al., 2001; Ketting et al., 2001; Knight and Bass, 2001). These primary siRNAs are loaded onto Argonaute proteins that bind to a target transcript through complementary base pairing, and then recruit RNA dependent RNA polymerases to promote formation of 22 nucelotide long secondary RNAs with a 5’ guanine (22G RNAs), which bear perfect complementarity to their targets. A distinct class of germline primary siRNAs termed piRNAs are 21 nucleotide small RNAs with a 5’ Uracil termed that are created from thousands of loci in the genome. piRNAs contribute to genome defence by silencing non-self transcripts such as transposons (Batista et al., 2008; Das et al., 2008; Shi et al., 2013; Wang and Reinke, 2008). piRNAs interact with the conserved Argonaute PIWI and can bind target RNAs with up to 3 mismatches, thereby triggering by recruitment of RDRPs and formation of 22G RNAs that are perfectly homologous to their targets and promote genomic silencing, similar to secondary siRNA formation in response to Dicer-dependent 26G primary siRNAs (Ruby et al., 2006; Shen et al., 2018; Zhang et al., 2018).

Consistent with a potential role of TERRA in telomere maintenance, siRNAs with homology to telomeres have been observed in a variety of organisms, where they may contribute to telomere maintenance. Small RNAs produced from telomere repeat-containing transcripts were found to promote DNA methylation at *Arabidopsis* telomeres (Vrbsky et al., 2010). Telomeric small RNAs have also been detected in mouse embryonic stem cells (Cao et al., 2009). Telomere dysfunction can induce production of telomeric siRNAs with perfect homology to the vertebrate telomere repeat sequence (TTAGGC)n, which are created by dsRNAs that arise from sites of DNA damage and are processsed by Dicer into siRNAs that promote the DNA damage response (Rossiello et al., 2017). However, the origin of small RNAs with homology to telomeres that are present under standard growth conditions of wildtype metazoan cells is not well understood.

Interstitial telomere sequences (ITSs) are degenerate telomeric sequences that are scattered along chromosome arms and are distinct from chromosome termini. A pioneering study by Meyne *et al.* identified cytologically visible ITSs in 55 different vertebrate species using fluorescence in situ hybridization (FISH) (Lin and Yan, 2008; Meyne et al., 1990). Azzalin *et al.* identified a number of ITS sites in the human genome by cloning methods and found that ITSs could be divided into three categories representing different putative mechanisms of origin: short, subtelomeric and fusion (Azzalin et al., 2001). Short ITSs are 24-130bp in length and display few mismatches with perfect telomere sequence. The authors suggested that these ITSs may be formed during DNA double-strand break repair. Subtelomeric ITSs comprise longer, more degenerate repeats that are found close to chromosome termini, and are likely to have originated from telomere recombination events. Fusion ITSs are likely a consequence of ancestral head-to-head telomere fusion events. Notably, only a single fusion-type ITS was identified in the human genome at 2q13, which led to the creation of chromosome 2 from two ancestral Hominid chromosomes (JW et al., 1991).

A number of studies have suggested a link between ITS sites and genome instability (Ashley and Ward, 1993; Bosco and de Lange, 2012; Kilburn et al., 2001; Lin and Yan, 2008). High mutation rates were observed at a region of telomeric sequence that was inserted into a yeast chromosome close to the left telomere (Aksenova et al., 2013; Moore et al., 2018). Intriguingly, the types of mutation observed depended on the orientation of the telomeric sequence. Insertion of the G-rich strand (with respect to the real left telomere on the chromosome) resulted in a high rate of indels and gross chromosomal translocations and inversions, while telomeric sequence inserted in the C-rich orientation experienced high rates of repeat loss and gain (Aksenova et al., 2015). Although ITSs can promote genomic instability, Shelterin components in mammals bind ITSs where they may act to stabilize these sequences by preventing the formation of secondary structures that interfere with DNA replication (Yang et al., 2011).

Inactivation of telomerase in *C. elegans* causes progressive sterility in the vast majority of animals. However, a small number manage to reproduce indefinitely in the absence of telomerase due to activation of ALT. In an analysis of stable *C. elegans* ALT strains, several strains were shown to have amplified a region of DNA between a telomere and an ITS sequence, which had then been trans-duplicated to the ends of the other chromosomes, where it acted as a surrogate telomere (Seo et al., 2015). This event is believed to have initiated from recombination between a critically shortened telomere and the ITS site, indicating that the high degree of similarity of ITS sites and telomeres has the potential to contribute to tumorogenesis and genome evolution by promoting a form of ALT-mediated telomere maintenance. However, despite the presence of ITSs in diverse species, the evolutionary origin and biological significance of these sequences remains mysterious.

Due to the combination of a wide availability of sequencing data and a genome that has been fully sequenced and assembled end-to-end, *C. elegans* provides a highly convenient model system in which to study small RNAs and genome biology. In this study, we analyzed published *C. elegans* small RNA-seq datasets and found numerous examples of small RNA species that map to perfect telomere sequence with up to three mismatches. We found that the majority of these small RNAs map to ITS sites, which are enriched for repressive chromatin marks and found mainly in the introns of protein-coding genes. We show that the closely related nematodes *C. briggsae* and *C. remanei* possess less abundant and generally more degenerate ITS sites than those in *C. elegans*, but display a substantially higher abundance of small RNAs that map perfectly to telomeres. We speculate that telomeric small RNAs may represent part of a functional pathway that has been significantly altered in *C. elegans* but retained in closely related species.

## Results

### Identification of small RNAs with high levels of telomere homology

We interrogated 75 publicly available small RNA sequencing libraries created from RNA isolated from various *C. elegans* strains and found rare reads that mapped perfectly to the *C. elegans* telomere repeat (TTAGGC)_n_, occurring at a frequency of 1 per 10,000,000 reads (Fischer et al., 2011; Gent et al., 2010; Gu et al., 2009; Phillips et al., 2015; Phillips et al., 2014; Sapetschnig et al., 2015; Seth et al., 2013) (Table S9). As piRNAs with up to 3 mismatches can target regions of the *C. elegans* genome for silencing (Bagijn et al., 2012), we counted telomeric siRNAs with up to 3 mismatches and found that these were substantially more abundant than those composed of perfect telomere repeat sequence (1 per 100,000) (FIg 1A). In total, 2352 distinct telomeric small RNA species were detected. Telomeric siRNAs with zero to two mismatches were significantly more likely to be composed of the telomeric G-strand sequence than would be expected by chance (p < 2.2e-16, binomial test) (Fig 1D). The proportion of telomeric siRNAs with one to three mismatches beginning with a G nucleotide was significantly greater than that for all siRNAs that could be mapped to the genome (p < 1e-15, Fisher’s exact test for all comparisons) and the median length for telomeric siRNAs with 1-3 mismatches RNAs was 22 nucleotides (Fig 1B,C). Therefore, telomeric siRNAs with 1-3 mismatches are enriched for the effector 22G RNA species, which has been established to promote either post-transcriptional or genomic silencing depending on the Argonaute they associate with (Billi et al., 2014). In contrast, rare telomeric siRNAs with 0 mismatches (perfect telomeric siRNAs) were not enriched for 22G RNAs but displayed peak lengths of 18 and 23 nt.

**Figure 1:**
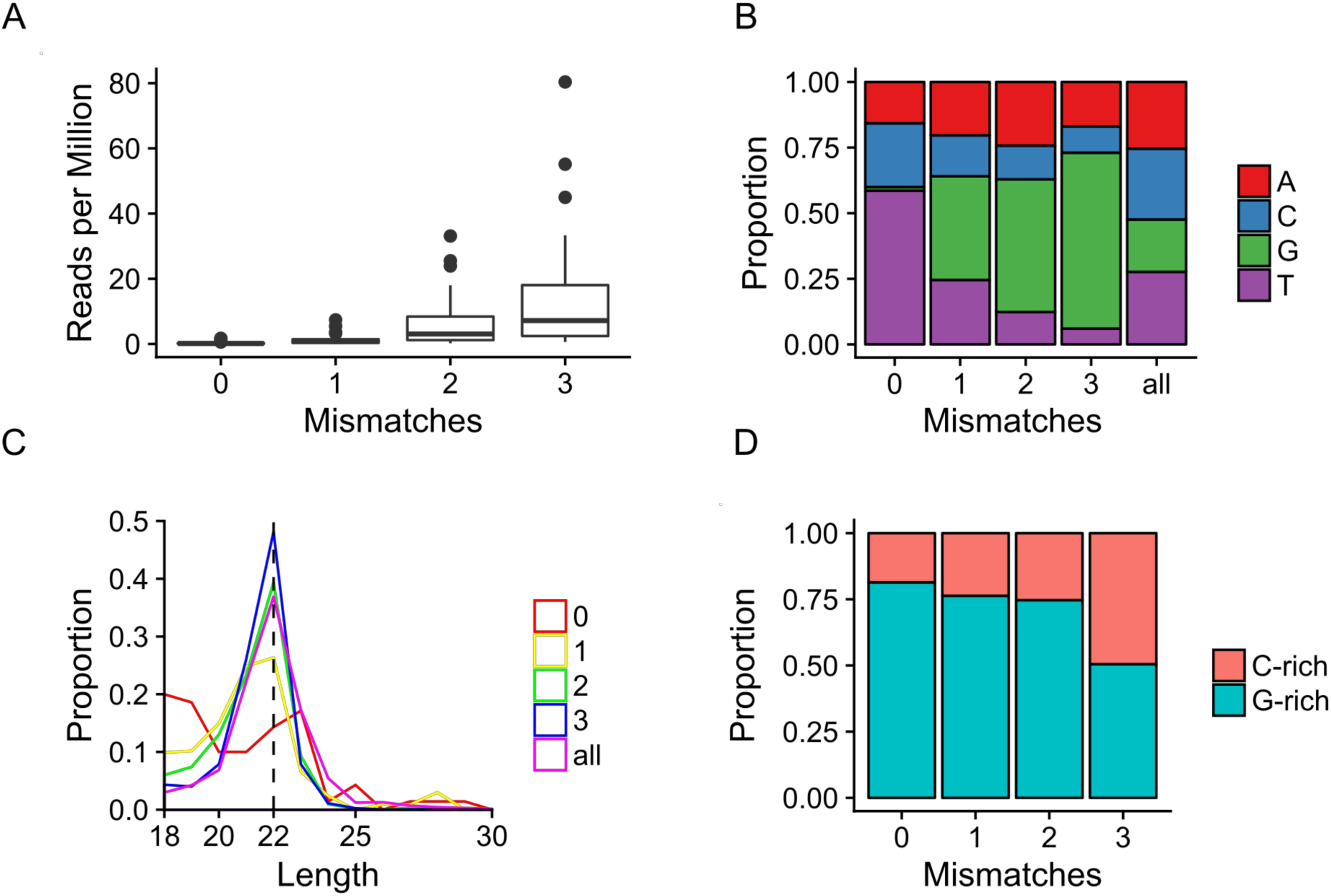
Identification of telomeric small RNAs in *C. elegans.* (A) Quantification of small RNA species in published datasets that map to telomere sequence with different numbers of mismatches. (B) Proportion of telomere-mapping small RNA reads in published data beginning with a particular nucleotide. (C) Distribution of telomeric small RNA length. (D) Proportion of telomeric small RNA reads mapping to the G-rich (TTAGGC) or C-rich (GCCTAA) telomeric strand. (E) Enrichment (IP/input) of reads mapping to perfect telomere sequence with up to three mismatches in published immunoprecipitation datasets for three different Argonaute proteins.

### ITS sites explain the origin of most telomeric small RNAs

Telomeric small RNAs could be divided into those that map to unique locations in the genome, which amounted to 10% and 57% of telomeric small RNAs with 2 or 3 mismatches, respectively, and whereas those that map to multiple locations, which amounted to 100% of telomeric small RNAs with 0 or 1 mismatches and 90% and 33% of telomeric small RNAs with 2 or 3 mismatches, respectively. The overall mapping frequency of telomeric small RNAs compares favorably with mapping frequency of all reads in a small RNA library, which generally amounts to 75-80% in our experience.

As unique telomeric small RNA reads mapped to loci that were scattered along chromosome arms, we reasoned that these loci may represent ITS sites that were previously shown to be concentrated on *C. elegans* chromosomes arms, based on the Ce000094 repeat sites that were automatically detected using the RECON algorithm (Bao and Eddy, 2002; Lowden et al., 2011) (https://wormbase.org/). Although the Ce000094 motif corresponds to telomeric DNA, we later noticed that there were many ITS sites that were covered only partially by this motif or not at all. This is likely a consequence of the degeneracy of ITS sites, which makes them challenging to detect de-novo using automated algorithms. In order to more thoroughly map ITS sites genome-wide, we located all instances of the telomeric hexanucleotide “TTAGGC” in the genome. We sought to capture perfect runs of telomere repeats as well as more degenerate sequences consisting of clusters of perfect repeats interspersed with non-telomeric sequence. TTAGGC hexanucleotides located on the same strand and within 100 nucleotides of each other were merged to form ITS contigs, and contigs that contained at least four perfect telomeric hexanucleotides were retained, resulting in 1229 ITS sites (Sup File 1).

ITS lengths ranged from 24 to 1204 nucleotides with a median length of 134 nucleotides (Fig 2F). ITS sites were found to be largely localized to the autosome arms, which have been shown to be largely heterochromatic and enriched for repetitive sequences (Fig 2A) (Garrigues et al., 2015). There was a fivefold depletion of ITS sites on the X chromosome (Fig 2B), consistent with lower levels of constitutive heterochromatin on this chromosome and an enrichment for Polycomb marks that silence the X chromosome in germ cells (Gaydos et al., 2012).

**Figure 2:**
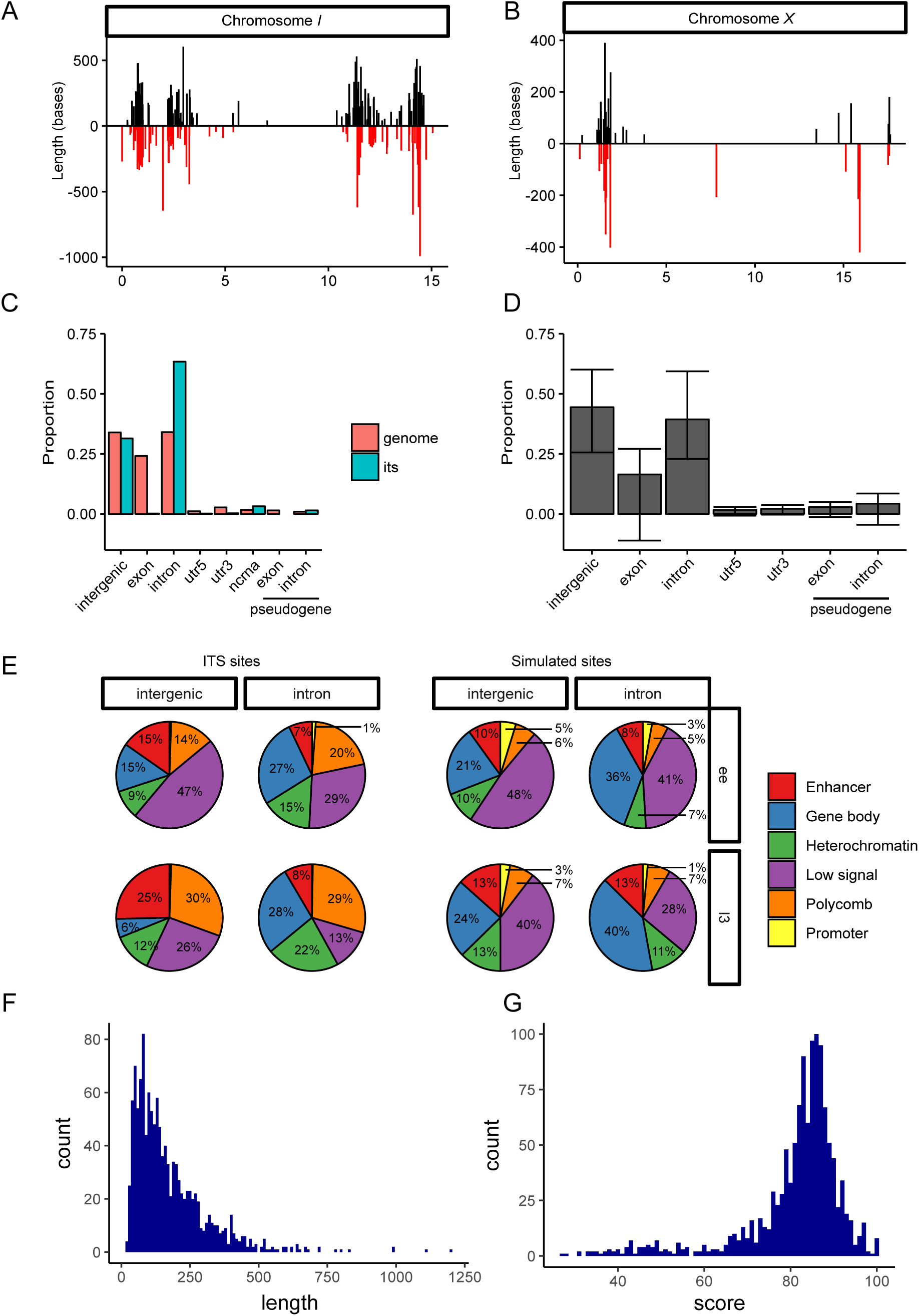
Characterization of ITSs in *C. elegans.* (A) Localization of ITS sites on chromosome I. regions in the sense orientation (TTAGGC) are shown in black, antisense in red. (B) Localization of ITS sites on the X chromosome. (C) Proportion of ITS sites overlapping different genome features (blue bars) and amount of genomic space taken up by genomic features (pink). (D) Average proportion of features overlapped by hexanucleotide repeat sequences from 1000 simulations. Error bars show standard deviation. (E) Percentage of real or permuted (“simulation”) ITS sites that are covered by a particular chromatin state. (F) Distribution of ITS length and score.

Although the mechanism by which ITS sites are created is unclear, these could be created by translocation of a telomere to an internal segment of the genome or by telomerase activity that acts at a double-strand break. Therefore, ITS sites are likely to be initially composed of perfect telomere repeats that degenerate over evolutionary time scales, such that they accumulate mutations in the absence of selective pressure. To obtain an estimate of the relative ages of ITS sites, we assigned a “homology score” to each site by performing local alignment between the ITS site and perfect telomere sequence of the same strand and length. The homology score was defined as the percent sequence identity between the two sequences. Homology scores ranged from 28.6 to 100, with a median score of 84.2 (Fig 2G, Sup File 1). Just over half (650) of ITSs had two or fewer consecutive perfect telomere repeats, although 25 ITS sites contained 7-15 consecutive perfect telomere repeats (Table 1). We found no obvious genome localization patterns in terms of homology score or sequence length. ITS sites therefore represent abundant repetitive sequences displaying a broad range of degeneracy (and therefore age) and genome localization patterns consistent with other types of repetitive sequence (Garrigues et al., 2015; Ikegami et al., 2010).

**Table 1:** Tally of the maximum number of consecutive perfect telomere repeats in ITS sites.

|  | Maximum consecutive perfect repeats |  |  |  |  |  |  |  |  |  |  |  |  |  |  |  |  |  |  |  |  |
| --- | --- | --- | --- | --- | --- | --- | --- | --- | --- | --- | --- | --- | --- | --- | --- | --- | --- | --- | --- | --- | --- |
| Species | 1 | 2 | 3 | 4 | 5 | 6 | 7 | 8 | 9 | 10 | 12 | 13 | 14 | 15 | 16 | 17 | 18 | 21 | 28 | 29 | 38 |
| *C. elegans* | 214 | 436 | 344 | 140 | 53 | 17 | 6 | 5 | 3 | 4 | 1 | 2 | 3 | 1 | 0 | 0 | 0 | 0 | 0 | 0 | 0 |
| *C. briggsae* | 344 | 15 | 0 | 0 | 2 | 2 | 1 | 1 | 0 | 2 | 1 | 0 | 1 | 0 | 0 | 1 | 1 | 0 | 0 | 0 | 0 |
| *C. remanei* | 164 | 99 | 7 | 0 | 1 | 2 | 2 | 1 | 1 | 1 | 1 | 0 | 1 | 1 | 1 | 1 | 1 | 1 | 1 | 1 | 1 |

**Table 2:** Summary of unique telomeric small RNA reads. Percentage of reads assigned mapping to ITS sites is shown.

| mismatches | reads | ITS | % ITS assignment |
| --- | --- | --- | --- |
| 0 | 29 | 23 | 79.3 |
| 1 | 182 | 167 | 91.8 |
| 2 | 769 | 640 | 83.2 |
| 3 | 1372 | 933 | 68.0 |

The origin of ITS sites is poorly understood. One proposed mechanism for ITS formation is the fusion of two chromosomes, although there currently exists evidence for only one such event in the human genome (JW et al., 1991). We sought to determine whether ancestral chromosome fusion events could account for the presence of ITSs in the *C. elegans* genome. However, we could find no example of an ITS site arranged head-to-head, and therefore conclude that ITS sites in the *C. elegans* genome are unlikely to arise as a result of chromosome fusions.

To determine whether ITS sites are a source of telomeric small RNAs, we mapped all small RNAs to the genome using a pipeline that assigns multi-mapping reads probabilistically based on the local density of uniquely mapped reads (Axtell, 2014). 76%, 91%, 83% and 68% of small RNAs mapping to telomeres with zero, one, two and three mismatches respectively were assigned to ITS sites (Table 1), whereas 92% and 75% of uniquely mapping small RNA reads with 2 or 3 mismatches respectively could be assigned to ITS sites. Although it is impossible to determine whether perfect telomeric small RNAs originate from telomeres or internal regions with certainty, these results suggest that such small RNAs are more likely to be produced from internal genomic regions. This is because many ITS sites are in heterochromatic segments of the genome and are surrounded by high densities of siRNAs, whereas few siRNAs map closely to subtelomeres. Furthermore, we looked for uniquely-mapping reads that overlapped with regions of perfect telomere repeat sequence of at least 30 bases in length, of which there were 95 in ITSs, and found 18 reads within internal regions, while only three reads could be found that overlapped telomere-subtelomere boundaries. As each ITS tract has two borders, this means that 18 small RNAs were present at these 190 ITS tract borders (1%), whereas 3 siRNAs were present at 12 telomere-sub-telomere boundaries (25%). These results suggest that perfect telomeric small RNAs could originate from either telomeres or ITS sites.

We asked whether telomeric small RNA abundance varied across ITS sites. Due to the uncertainties associated with multi-mapping reads, we decided to focus on the uniquely-mapping subset of telomeric small RNAs with two or three mismatches. We could find no strong correlation between telomere homology score and small RNA mapping frequency for ITS (Spearman’s ρ = 0.068), although the ITS sites with the highest telomeric small RNA abundance had homology scores of around 80% and no telomeric small RNAs were found at sites with scores less than 53.8% (Fig 3A). For example, all ITS tracts with >10 telomeric small RNAs had homology scores of >76% (Table 3). The abundance of uniquely mapping telomeric small RNAs correlated well with total small RNA abundance at an ITS tract (Spearman’s ρ = 0.70), indicating that ITS sites that produce high numbers of telomeric small RNAs are likely to be sites of high small RNA production in general. Of the top 10 ITSs with the highest overall small RNA abundance, seven also displayed high telomeric small RNA abundance. The remaining three ITSs that had no telomeric small RNAs assigned had homology scores of less than 56%. We found one ITS site (ITS_60) which displayed an unusually high frequency of small RNA mapping, 5- to 13-fold greater than other ITS tracts with abundant siRNAs and accounting for 43% and 18% of total telomeric small RNAs with two and three mismatches respectively (Fig 3, Table 3). This ITS is intergenic and fully overlaps two annotated converging non-coding RNAs, whose transcripts could anneal to produce dsRNA that can be processed into abundant ITS siRNAs. (Fig 3B). As telomeric small RNAs did not map exclusively to ITS sites, we also looked at the genome-wide distribution of telomeric small RNAs (Fig 3C). We found three peaks of high coverage that did not overlap any ITS sites. Two of these, located on the left arms of chromosomes III and V, contained two consecutive perfect telomeric repeats and were assigned a number of reads with two or three mismatches to telomeres, but none with zero or one. The other peak, located on the right arm of chromosome I, contained three consecutive perfect telomere repeats next to a degenerate one, narrowly missing the criteria for being detected as an ITS site. Therefore, telomeric small RNAs that do not originate from ITS sites are likely to originate from non-ITS regions which happen to have short stretches of telomeric DNA.

**Table 3:** ITS sites with more than 10 telomeric small RNAs.

| ID | chromosome | start | end | length | score | type | telomeric small RNAs |  |  |
| --- | --- | --- | --- | --- | --- | --- | --- | --- | --- |
|  |  |  |  |  |  |  | 2 | 3 | all |
| ITS_60 | I | 1974471 | 1975116 | 645 | 81.9 | intergenic | 30 | 105 | 135 |
| ITS_813 | IV | 1752185 | 1752299 | 114 | 86.8 | intergenic | 5 | 22 | 27 |
| ITS_238 | I | 14733054 | 14733310 | 256 | 78.7 | intron | 0 | 22 | 22 |
| ITS_1226 | X | 17527653 | 17527730 | 77 | 75.6 | 3' UTR | 0 | 22 | 22 |
| ITS_576 | III | 1881503 | 1881935 | 432 | 82.9 | intron | 2 | 19 | 21 |
| ITS_544 | III | 953856 | 953988 | 132 | 76.1 | intergenic | 0 | 18 | 18 |
| ITS_316 | II | 2859211 | 2859720 | 509 | 79.5 | intergenic | 3 | 13 | 16 |
| ITS_739 | III | 12856138 | 12856366 | 228 | 79.4 | intron | 0 | 16 | 16 |
| ITS_315 | II | 2847764 | 2848131 | 367 | 79.6 | intron | 2 | 13 | 15 |
| ITS_314 | II | 2846566 | 2846973 | 407 | 76.4 | intron | 0 | 11 | 11 |

**Figure 3:**
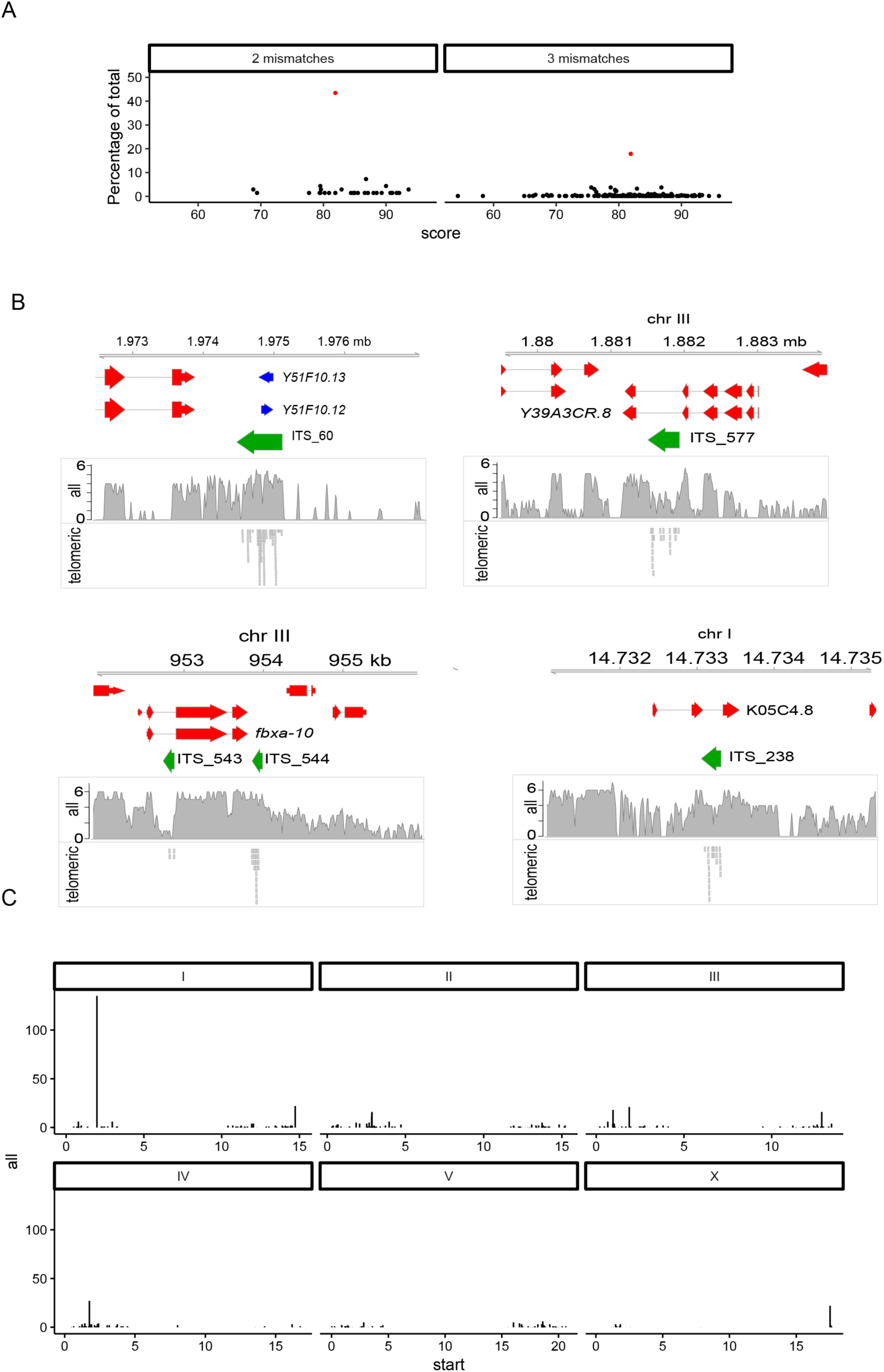
Telomeric small RNAs map to ITS sites. () Scatter plots showing ITS score versus uniquely-mapping small telomeric small RNAs count as a percentage of total abundance for each category for ITS sites with at least one small RNA read. ITS_60 is shown in red. (B) Genome track showing the top four ITSs with the highest uniquely-mapping telomeric small RNA count. Total small RNA coverage and telomeric small RNA reads are shown as separate tracks (C) Genome-wide distribution of ITS sites. Y-axis shows uniquely-binding telomeric small RNA count.

### ITS sites are mainly found in introns and are enriched for repressive chromatin marks

We looked for overlap between ITS sites and annotated genomic features and found that they were enriched in introns and in intergenic sequences. Surprisingly, 796 ITS sites (63% of total) were located in introns of protein coding genes (Fig 2C). In terms of sequence coverage, this represents almost a two-fold enrichment for ITS sequence at introns based on the proportion of the genome that is comprised of intronic sequence. In contrast, only two ITS sites (0.16%) were found in protein-coding exons, indicating that ITS sites are depleted from exome, possibly because they are deleterious for mRNA expression. Consistently, ITS sites were also significantly depleted from both 5’ and 3’ UTRs. Such a pattern could be a general feature of blocks of DNA comprised largely of degenerate hexanucleotide repeats. To test this, we repeated the ITS detection pipeline with 100 random hexanucleotide combinations. For each hexamer, we defined degenerate repeat sites using the same method used to detect ITS sites and looked for overlap between these sites and annotated genomic features. The simulated repeat sites did not show an overall enrichment at introns or depletion at exons (Fig 2D), indicating that this pattern of localization is specific to ITS sites. Genes containing intronic ITS sites were not significantly enriched for any GO terms.

Features of eukaryotic chromatin such as histone modifications and nucleosome occupancy drive spatio-temporal regulation of genomic regions. Ho *et al.* employed a hidden Markov model based algorithm to classify chromatin into a series of “states” based on the integration of various CHIP-seq and microarray datasets (Ho et al., 2014). The 16 states were grouped into six categories: promoter, enhancer, gene body, Polycomb-repressed, heterochromatin, and weak or low signal. We assigned chromatin states to each ITS sequence using the *C. elegans* dataset. To determine statistically significant enrichment for chromatin states, we performed 1000 simulations in which a new set of 1219 sequences with the same length distribution as the ITS sequences was randomly selected from the genome and assigned chromatin states. Both intronic and intergenic ITS sites were enriched two to four-fold for Polycomb-repressed chromatin states in both the early embryo and at the L3 larval stage of development relative to the corresponding simulated sites (p < 0.0003 for all classes), with the highest enrichment in the L3 stage (Fig 2E, Table S1). Intronic but not intergenic ITS sites were enriched two-fold for heterochromatin chromatin states (p = 0.008 and 0.02 for early embryo and L3 respectively) and were depleted for enhancer chromatin states. The chromatin state of intronic ITSs matched that of the nearest exon in over 99% of cases for both early embryo and L3, indicating that the chromatin environment of intronic ITSs is almost always shared by the entire gene. Intergenic ITS sites were enriched 1.5-fold and two-fold for enhancer states in early embryo and L3 larvae respectively (p < 0.00048, p = 0.0003 for L3 larvae and early embryo respectively). We looked at the genes most proximal to intergenic ITSs with enhancer chromatin states but found no significantly enriched GO terms amongst these genes. Both intronic and intergenic ITS sites were strongly depleted for promoter chromatin states at both stages (p < 0.001 in each case). We found no significant relationship between chromatin state and proportion of ITSs that produce at least five small RNAs (p = 0.1, Chi Square test for both early embryo and L3 stages). We conclude that ITS sites are generally associated with repressive chromatin, that intronic ITS sites may promote repressive chromatin formation via heterochromatin or Polycomb pathways, but that some intergenic ITS sites may possess enhancer function.

### *C. briggsae* and *C. remanei* ITS sites are less common and more degenerate than those of *C. elegans*

We sought to gain insights into the evolutionary conservation of ITS sites and the relationship between ITS sites and small RNA production in nematodes. *C. briggsae* is one of the closest known living relatives of *C. elegans*, and diverged from *C. elegans* between 18-100 million years ago (Cutter, 2008; Stein et al., 2003). It is physically virtually indistinguishable from *C. elegans* and similarly reproduces through selfing. *C. remanei* is another close relative of *C. elegans*. Although *C. briggsae* and *C. remanei* may have shared a recent common ancestor that diverged from *C. elegans, C. remanei* reproduces by gonochoristic male-female sexual reproduction, whereas *C. elegans* and *C. briggsae* are both hermaphroditic selfers (Fierst et al., 2015).

We applied the same method for detecting ITS sites in *C. briggsae* and *C. remanei* and found only 372 and 292 sites respectively, less than one third of the number found in *C. elegans* in each case. ITS sites in *C. briggsae and C. remanei* were significantly more divergent than those from *C. elegans*, with median homology scores of 49.5 and 57.1 (p < 2.2e-16, Mann-Whitney U test for both comparisons) (Fig 4A), while the median length for ITS sites was slightly higher for *C. briggsae* and *C. remanei* at 206 (p < 2.2e-16, Mann-Whitney U test) and 178 nucleotides (p = 6.058e-09, Mann-Whitney U test), respectively (Fig 4B). Interestingly, although 93% and 57% of ITSs in *C. briggsae* and *C. remanei* contained no consecutive telomere repeats, both species possessed a handful of 36-228 nucleotide ITSs with perfect telomere homology (Table 1, Sup file 1). Although *C. remanei* had the longest perfect telomere tracts of the three species, *C. remanei* and *C. briggsae* possessed few ITS tracts with long stretches of perfect telomere repeats in comparison to *C. elegans* (Table 1). All three nematode genomes showed an intron bias for ITS sites, which in comparison to *C. elegans* was stronger for *C. briggase* and weaker for *C. remanei.* (Fig 4C, D). It was not possible to determine an enhancer bias, because the other genomes have not been annotated for this feature.

**Figure 4:**
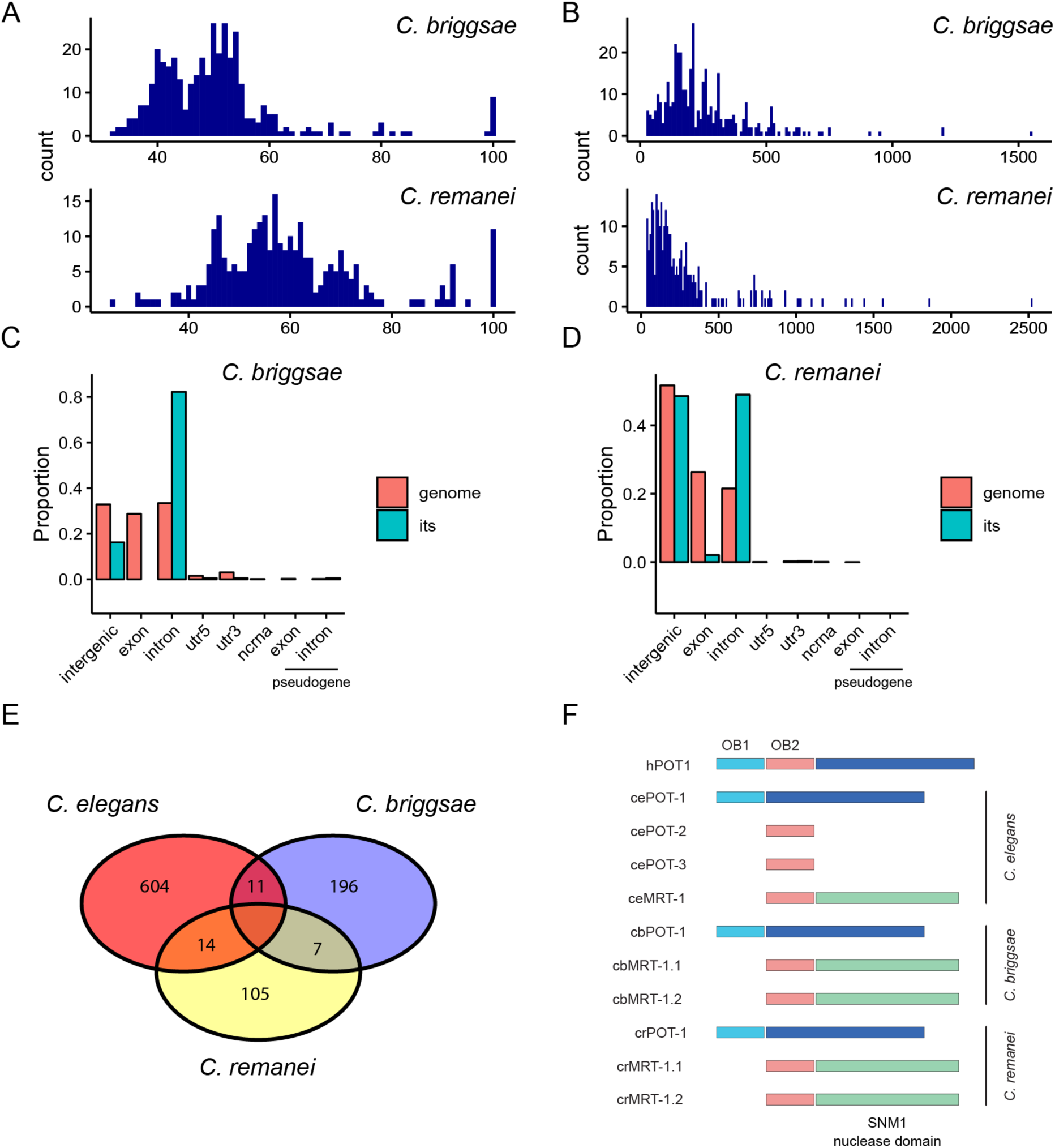
ITS sites in *C. briggsae* and *C. remanei*. (A) Distribution of ITS scores. (B) Distribution of ITS lengths. (C) Proportion of ITS sites overlapping different genome features (blue bars) and amount of genomic space taken up by genomic features (pink). (D) As (C) but for *C. remanei*. (E) Number of genes for which orthologous genes in two species containin intronic ITS sites in both species. (F) Domain structures of *C. elegans, C. briggsae and C. remanei pot-2* homologues

To gain insights into the evolutionary history of ITSs across different species, we looked for intronic ITSs that are shared within orthologous genes. We could not identify genes for which orthologues between all three species contained ITS sites. We performed pairwise comparisons between the three nemotode species and found a handful of genes in each case for which orthologous genes between the two species contained ITS sites in both species, which we termed “shared” ITS sites as opposed to “unique” sites (Fig 4E, Table S2). We found that the median score was slightly but significantly lower for shared sites compared with unique sites in C. *elegans* (78.7 versus 84, p = 0.008443, Mann-Whitney U test), although no significant difference was noted for *C. briggsae* or *C. remanei* (Figure S1). Notably, there were no shared ITS sites with scores greater than 94.4%, consistant with unique ITS sites having been formed more recently in evolution. We performed GO term analysis on *C. elegans* genes orthologous of *C. briggsae* and *C. remanei* genes that contain ITSs. The *C. briggsae* gene set was enriched for GO terms related to reproduction cell cycle and organelle organization. No significant GO terms were enriched in the *C. remanei* gene set. Therefore, the mechanism of ITS formation is ancient in origin but ITSs themselves are divergent among nematode species, suggesting that a possible function of ITS tracts in regulation of chromatin if present diverges rapidly between species.

### Telomeric small RNAs are more abundant in *C. elegans* relatives

We then interogated small RNA libraries in *C. briggsae* and *C. elegans* (Shi et al., 2013) and looked for small RNAs that map to telomeres. The same dataset contained a *C. elegans* library, which we also analyzed as a control. Consistent with our previous analysis, reads mapping perfectly to telomeres were exceedingly rare in the *C. elegans* libraries; three were detected in hermaphrodites, one in embryos and none in males, corresponding to a total of around one read per 10,000,000 small RNA reads. Strikingly, libraries from *C. briggsae* and *C. remanei* contained several orders of magnitude more perfect telomeric small RNAs compared with *C. elegans* (Fig 5A, Sup file 2). Perfect telomeric small RNAs were detected at an abundance of more than 22 per million in *C. briggsae* embryos and mixed male/female *C. remanei* adults. The *C. elegans* perfect telomeric small RNAs detected in control libraries from the *briggsae* and *remanei* datasets showed the same G-rich strand bias as found in the libraries described earlier. However, the *C. briggsae* and *C. remanei* libraries showed a strong bias for the C-rich strand bias for siRNAs composed perfect telomere repeats (Fig 5B). Despite this strong distinction in strand bias, perfect telomeric small RNAs from all three species showed a striking depletion of perfect telomeric small RNAs species that begin with a 5’ G, whereas abundant levels of small RNAs with a 5’ guanine were observed for telomeric siRNAs with mismatches.

**Figure 5:**
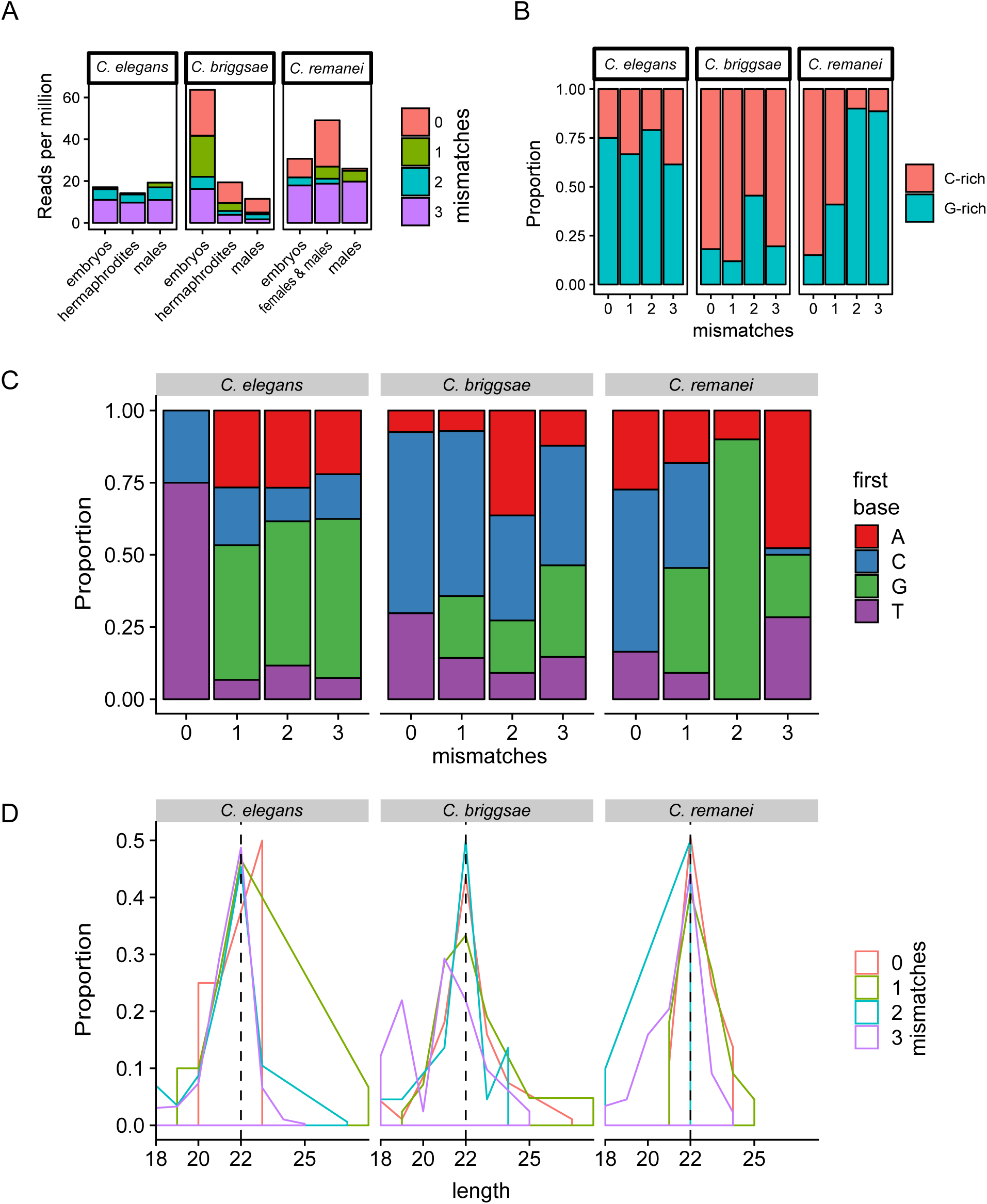
Comparison of telomeric small RNAs between three *Caenorhabditis* species. (A) Telomeric small RNA abundance in small RNA sequencing libraries (REF) from three *Caenorhabditis* species. (B) Proportion of telomeric small RNA reads mapping to the G-rich (TTAGGC) or C-rich (GCCTAA) telomeric strand. (C) Proportion of telomere-mapping small RNA reads in published data beginning with a particular nucleotide. (D) Distribution of telomeric small RNA length.

If the perfect telomeric siRNAs are split into G- and C-strand categories, the G-rich RNAs from all three species were enriched for RNAs beginning with 5’ UAGGCU or 5’UUAGGC, ranging in size from 18-29 nucleotides. The most abundant sRNAs from all three species were 23 nt RNA UUAGGCUUAGGCUUAGGCUUAGG and 22nt RNA UAGGCUUAGGCUUAGGCUUAGG (Fig 6). On the other hand, C-rich telomeric RNAs of *C. briggsae* and *C. remanei* species showed enrichment for 22 nucleotide perfect telomeric small RNAs beginning with a C (Fig. 5C,D) of 30% and 32%, respectively. Most perfect C-rich telomeric RNAs both *C. briggsae* and *C. remanei* began with 5’ CUAAGC and ranged from 18-24 nt in length, with the most abundant C-rich RNA being the 22 nucleotide sequence CUAAGCCUAAGCCUAAGCCUAA, which comprised 22% and 29% of all sequences for *C. briggsae* and *C. remanei* respectively (Table S2). This sequence reflects the majority of 22 nt perfect telomeric small RNAs for each species (52% and 57%, respectively). Given that there are only six permutations of a perfect telomeric sequence of a given length, each permutation would be expected to occur at a probability of 1/6 if the sequence distribution was random. We conclude that the general depletion of 5’G nucleotides from perfect telomeric siRNAs can be largely explained by the fact that G-rich RNAs commonly begin with 5’UAGGCU and 5’ UUAGGC, whereas C-rich RNAs commonly begin with 5’CUAAGC (Fig. 6 and Table S2).

**Figure 6:**
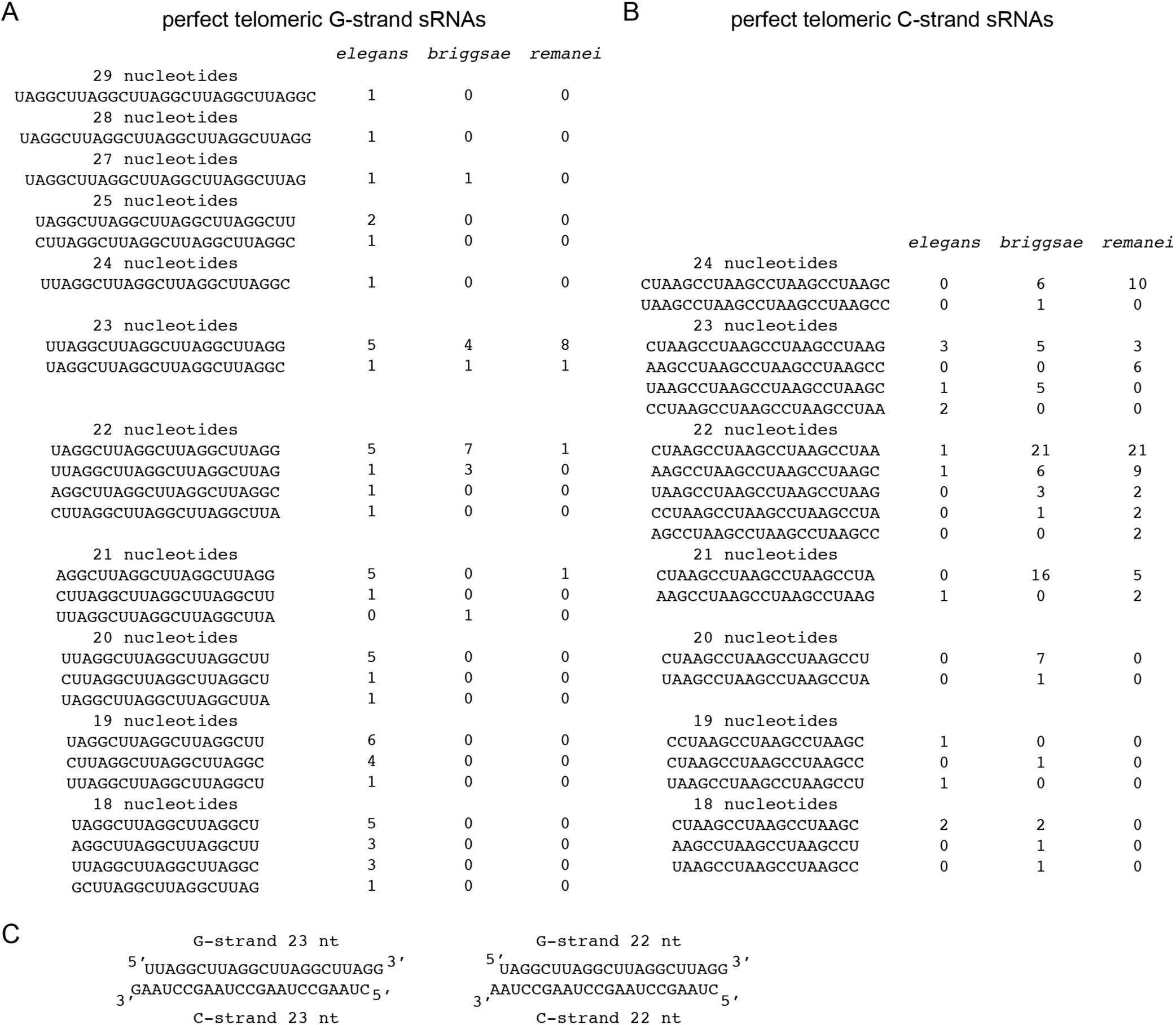
Perfect telomeric small RNAs of the genus *Caenorhabditis*. Total observed G-rich (A) and C-rich (B) telomeric small RNAs from RNA seq and IP data sets. (C) Abundant telomeric sRNAs can anneal to form duplexes.

Due to the relatively small number of available small RNA libraries for *C. briggsae* and *C. remanei* and the fact that telomeres have not been assembled in the reference genomes for these species, we could not make inferences about whether telomeric small RNAs were more likely to originate from telomeres or ITSs based on siRNAs that map to borders of perfect telomere repeat tracts. If perfect telomeric small RNAs originate from ITS sites, an increase in abundance for these RNAs could be due to an increase in the amount of ITS sequence that is capable of producing them. However, the *C. elegans genome* contains a combined total of 3666 bases of perfect telomere existing in stretches of at least 30 nucleotides in length, compared with 1596 for *C. briggsae* and 708 for *C. remanei*, indicating that increased levels of perfect telomeric ITS sequence cannot account for the increase in perfect telomeric small RNA abundance. We conclude that the composition of telomeric small RNAs in *C. elegans* has diverged substantially from its close relatives, but that both classes of small RNAs are dramatically depleted for 5’ G RNA species, which could reflect either a toxicity of this species or its mechanism of biogenesis.

Argonaute proteins can promote the biogenesis of specific classes of small RNAs. For example, ALG-3 and ALG-4 associate with 26G RNAs and PRG-1/Piwi associate with 21U RNAs, whose levels are dramatically reduced in response to deficiency for *prg-1* (Batista et al., 2008; Das et al., 2008; Han et al., 2009; Ruby et al., 2006; Vasale et al., 2010). We studied small RNAs that immunoprecipitate with Argonaute proteins PRG-1, WAGO-1, HRDE-1 and CSR-1 (Ashe et al., 2012; Batista et al., 2008; Buckley et al., 2012; Claycomb et al., 2009; Das et al., 2008; Gu et al., 2009). Perfect telomeric small RNAs were exceedingly rare in the input RNA datasets, as expected, and we found no enrichment for C- or G-strand telomeric RNAs in any IP dataset (Table S3). However, these datasets revealed the only three *C. elegans* examples that we observed of the major C-strand RNA species found in *C. briggsae* and *C. remanei*, 23 nt 5’ CUAAGC RNA. The IP datasets brought the total number of *C. elegans* C-rich telomeric RNAs to 14 (Fig. 6, Tables S2 and S3). Significantly numbers of telomeric small RNAs with mismatches were associated with PRG-1, WAGO-1, HRDE-1 and CSR-1 Argonaute proteins.

### Evolution of telomeric small RNAs

Perfect telomeric siRNAs have been previously reported to associate with the *Tetrahymena* Argonaute protein Twi10, were of lengths 23-24nt and 33-36nt and all possessed the sequence 5’UGGGGU (Couvillion et al., 2009), potentially similar to the 5’ UAGGCU G-rich RNAs that were present in *Caenorhabditis* species. Telomeric siRNAs have been previously reported to be rare in mammals, where they may promote the response to DNA damage at telomeres. We therefore examined ∼650,000,000 reads in 77 small RNA libraries from mouse and found 156, 56, 88 and 860 telomeric reads with 0, 1, 2 and 3 mismatches, respectively (Tables S5-S6,S9) (Manakov et al., 2015; Pandey et al., 2013; Watanabe et al., 2015; Wenda et al., 2017). Perfect telomeric siRNAs occurred at a frequency of ∼1 in 4,170,000 reads and were therefore almost as rare of *C. elegans* perfect telomeric siRNAs. G-rich perfect telomeric siRNA were also more common in mammalian telomeric siRNAs with the most abundant beginning with 5’ GUUAGG and were 19 and 23 nt in length (Tables S7). The most common C-rich telomeric siRNA species from mammals begin with 5’UAACCC and were 24, 28 and 30 nt in length (Table S8). Other permutations of 5’ ends were represented for both G- and C-rich perfect telomeric RNAs from mammals.

We speculated that the striking difference in telomeric small RNA composition between *C. elegans* and its relatives could occur in response to a functional difference in telomere biology between these species. The *C. elegans* shelterin complex contains the OB-fold containing proteins POT-1 and POT-2, and mutation of either *pot-1* or *pot-2* results in successive telomere elongation over multiple generations (Raices et al., 2008; Shtessel et al., 2013). Variants in the *pot-2* gene have also been found to have a prominent effect on telomere length in wild *C. elegans* strains, suggesting that this protein may affect fitness in the wild (Cook et al., 2016). A third *C. elegans* homolog of human Pot1, MRT-1,is required for telomerase activity (Cook et al., 2016). We performed tBLASTX searches for orthologues of the POT-2 protein in *C. briggsae and C. remanei* genomes, respectively. Surprisingly, we found that the reference genomes of *C. briggsae* and *C. remanei* do not contain *pot-2* homologues. Instead, these species each posses two copies of MRT-1, which has an SNM1DNA interstrand crosslink repair nuclease fused to the POT-2 OB-fold protein (FIg. 4F) (Meier et al., 2009). It is possible that the absence of *pot-2* or presence of an extra *mrt-1* paralogue in *C. briggsae* and *C. remanei* could represent a fundamental difference in telomere biology that is related to the increased abundance of perfect telomeric small RNAs in these species.

We examined telomeres of several wild isolates of *C. briggsae* and *C. remanei* and found that the AF16 strain that has high levels of perfect telomeric siRNAs had elongated telomeres, whereas the other strains examined had generally short telomeres, similar in size to those of *C. elegans*. However, several *C. briggsae* and *C. remanei* strains possessed weak telomeric DNA signals that failed to migrate into the gel and that ran at limit mobility, which are not observed in the genomic DNA from the Bristol N2 wildtype *C. elegans* strain, but commonly occur in genomic DNA from *C. elegans* strains of *trt-1* mutants whose telomeres are maintained by the telomerase-independent telomere maintenance pathway ALT (Fig. 7) (Cheng et al., 2012).

**Figure 7:**
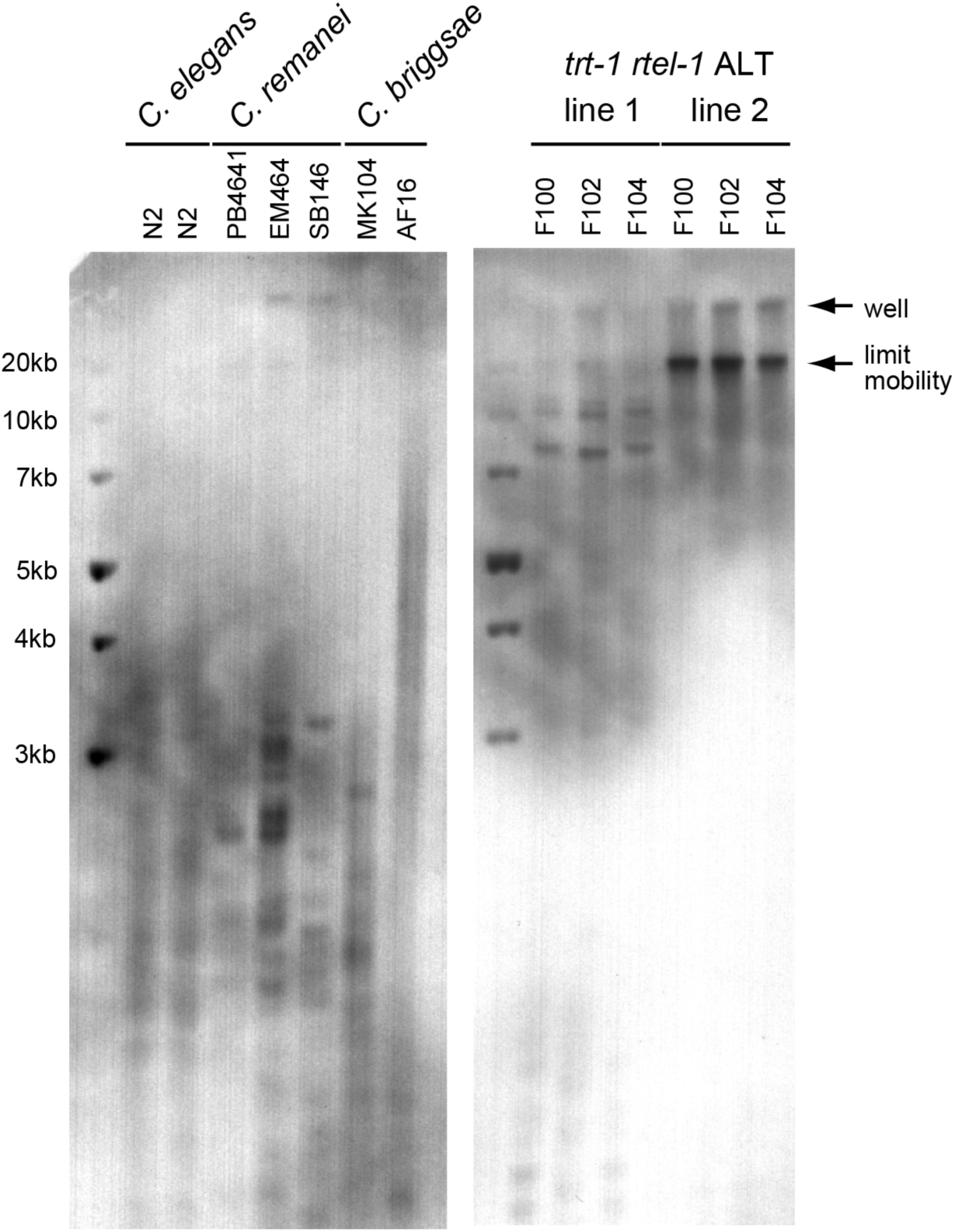
Telomere length of *C. elegans, C. briggsae* and *C. remanei* strains. (A) Southern blot analysis of Hinf1-digested genomic DNA isolated from *C. elegans, C. briggsae* and *C. remanei* strains using a (TTAGGC)_n_ probe. (B) Southern blot analysis of two *C. elegans trt-1* telomerase mutant strains whose telomeres are maintained via ALT. Positions of telomeric bands located in the well or at limit mobility are noted by arrows.

## Discussion

Small RNAs have been shown to occur at telomeres in several different organisms (Couvillion et al., 2009; Rossiello et al., 2017; Vrbsky et al., 2010). We interrogated a number of published *C. elegans* small RNA sequencing datasets and found rare examples of small RNAs mapping perfectly to telomeres as well as more abundant species mapping to telomeres with up to three mismatches. Telomeric small RNAs with an increasing number of mismatches correlated with an increasing enrichment for 22G small RNA species, which represent secondary ‘effector’ siRNA molecules that are produced de novo by RNA-dependent RNA polymerases during both exogenous and endogenous RNA inteference. All telomeric small RNAs mapping to telomeres with zero or one mismatches and the majority of species with two or three mismatches could be mapped to ITS sites using a method that randomly distributes non-unique reads, suggesting that ITS sites could be responsible for the formation of perfect telomeric small RNAs. However, mapping of siRNAs to borders of ITS tracts and of telomeres suggested that while long stretches of perfect telomere repeats at either ITS tracts or at telomeres can give rise to siRNAs, telomeres could represent the more abundant source of perfect telomeric small RNAs. Perfect telomeric small RNAs that are created from ITS tracts might possess characteristics of small RNAs derived from ITS tracts, such as beginning with any 5’ nucleotide, which could explain the range of 5’ nucleotides that were observed for perfect G- or C-strand telomeric small RNAs.

Strikingly, perfect telomeric small RNAs were substantially more abundant in *C. briggsae* and *C. remanei*, whereas telomeric small RNAs with mismatches were generally of comparable abundance to *C. elegans.* However, perfect telomere repeat sequences within ITSs were depleted in *C. briggsae* and *C. remanei* relative to *C. elegans* (Table 1), suggesting that telomeres rather than perfect telomere sequence within ITS tracts are likely to be the source of the majority of perfect telomeric small RNAs. In parallel with their altered level of abundance, perfect telomeric small RNAs of *C. briggsae* and *C. remanei* displayed a substantial bias for being composed of the C-rich strand, which contrasts from the G-rich bias of small RNAs from *C. elegans*. The overrepresentation of C-rich telomeric small RNAs in *C. briggsae* and *C. remanei* is puzzling, possibly reflecting a change to the mechanism of telomeric small RNA biogenesis. In this regard, the C-rich telomere small RNA species would be complementary to the telomeric non-coding RNA TERRA, which is considered to be the most abundant telomeric RNA (Azzalin and Lingner, 2015). We note that mammalian telomeric small RNAs were more frequently composed of the G-rich telomeric sequence and that their levels were very low, as observed for *C. elegans.* Together, our results argue that small RNA classes that are very rare may be functionally significant, which is consistent with a proposed role for telomeric small RNAs in responding to DNA damage at mammalian telomeres (Rossiello et al., 2017).

Perfect telomeric G-rich RNAs from all three *Caenorhabditis* species commonly began with 5’UAGGCU and 5’ UUAGGC. C-rich telomeric small RNAs of *C. briggsae* and *C. remanei* were enriched for 5’ ends that began with 5’CUAAGC, which occurred at 6 of 14 in *C. elegans* C-rich RNAs (Fig. 6). Despite having prominent 5’ end sequences, both G- and C-rich telomeric RNAs displayed a wide range of lengths, suggesting that they may have a common mechanism of biogenesis that may be distinct from other known *C. elegans* pathways that often display pronounced enrichment for a discrete RNA length. Perhaps consistently, *Tetrahymena* possesses telomeric small RNAs with 5’ UGGGGU ends that also displayed a range of sizes (Couvillion et al., 2009). While we did not observe consistent 5’ sequences for mammalian perfect telomeric small RNAs, these were biased towards the G-rich telomeric siRNAs and were extremely rare, as observed for *C. elegans* telomeric small RNAs. We found that 3’ ends for major species of G-rich small RNAs for *Caenorhabditis* species and mammals aligned perfectly with the 5’ end of the C-rich DNA base at the chromosome terminus (Fig. 8).

**Figure 8:**
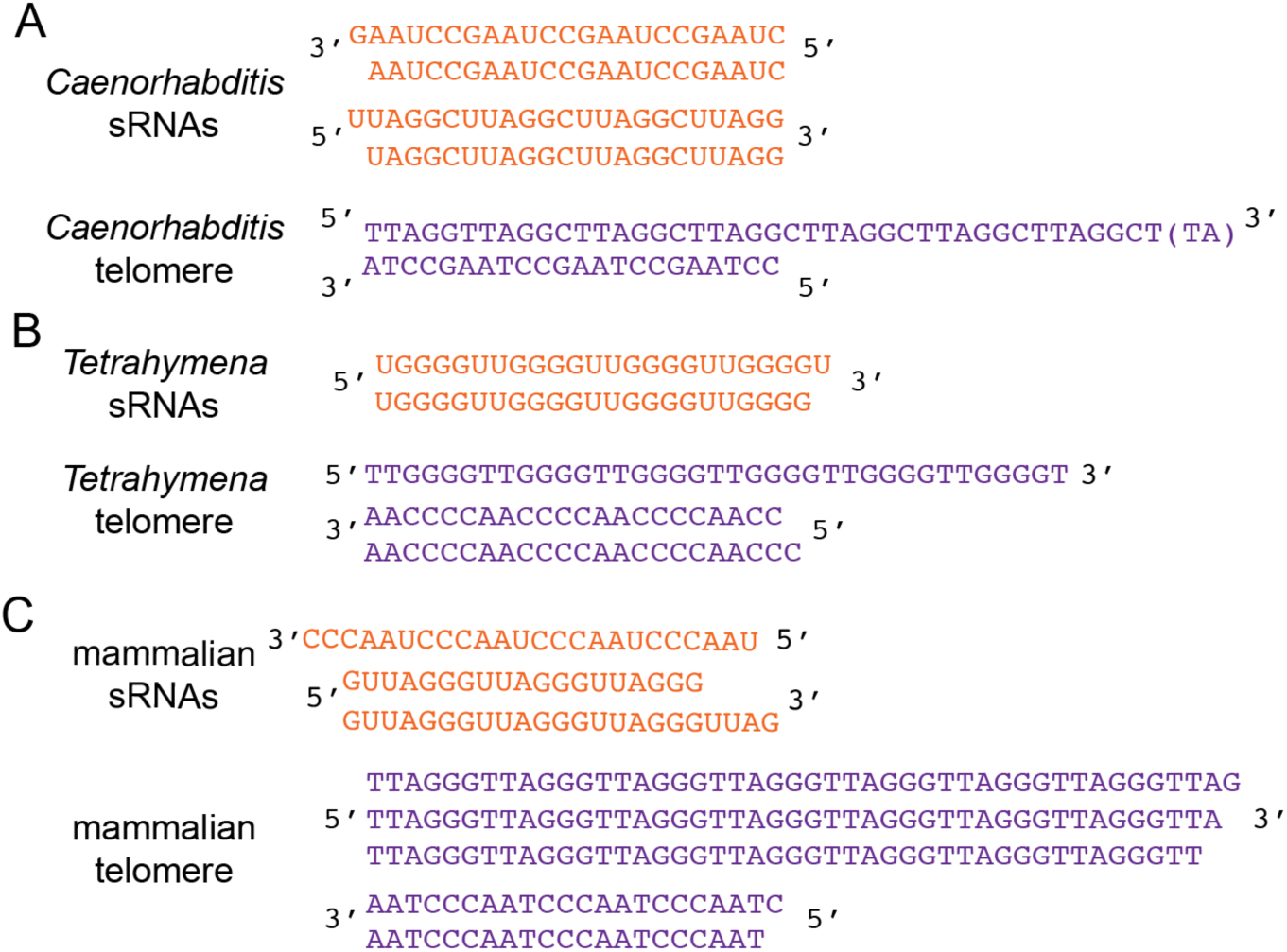
Models of telomere RNAs and chromosome termini. Comparisons of prominent species of small RNAs (orange) and chromosome termini (lavender) for (A) *Caenorhabditis* (Raices et al., 2008) (B) *Tetrahymena* (Jacob et al., 2003; Jacob et al., 2001), and (C) mammals (Sfeir et al., 2005). The *Caenorhabditis* terminal (TA) dinucleotide indicates uncertain nucleotides at the end of the 3’ overhang, whereas the *Tetrahymena* and mammalian 3’ nucleotides have been confirmed experimentally, the most frequent indicated based on highest vertical position.

Given that *C. briggsae* and *C. remanei* are more closely related to each other than to *C. elegans*, we examined their genomes and found consistent loss of the POT-2 protein that represses telomerase activity and ALT in *C. elegans*. Some wild isolates of *C. elegans* have long telomeres (Raices et al., 2005), and this is sometimes due to a loss of function mutation in *pot-2* (Cook et al., 2016). We examined telomeric DNA from several wild isolates of *C. briggsae* and *C. remanei* and found it to be distinct from Bristol N2 *C. elegans* DNA in that minor fraction of their telomeres adopted were long or unusually structured, similar to what is observed in telomerase-negative strains where ALT is active. Together, there results suggest that either loss of *pot-2* or high levels of telomeric small RNAs is associated with ALT-like telomeric DNA observed in *C. briggsae* and *C. remanei* strains. It is possible that loss of POT-2 in *C. elegans* promotes turnover of telomeric small RNAs or that the second MRT-1 homolog of *C. briggsae* and *C. remanei* promotes biogenesis of telomeric siRNAs. Both POT-2 and MRT-1 have OB2-folds that are predicted to interact with 3’ ends of single-stranded telomeric DNA (Lei et al., 2003), and Argonaute proteins possesses a PAZ domain that uses a variant OB fold to interact with the 3’ termini of single-stranded RNA (Song et al., 2003).

We found that ITS sites are a shared feature amongst close relatives of *C. elegans*, although their compositions bare some striking differences. ITSs in both *C. briggsae* and *C. remanei* are rarer and generally more degenerate than in *C. elegans*, although both species contain examples of more recently created ITS sites with long stretches of perfect telomere homology. It is intriguing that there should be such a discrepancy in ITS divergence between these species, and this observation suggests ITS tract creation may be under selective pressures. Although it may be a coincidence that perfect telomeric small RNAs are far more abundant in *C. briggsae* and C. *remanei*, it is possible that these RNAs are functionally linked to ITS formation and/or evolution. For example, perfect telomeric small RNAs may actively inhibit formation or expansion of ITS sites, meaning that their reduced levels in *C. elegans* has led to an altered landscape of ITSs in the *C. elegans* genome, many of which have been formed relatively recently.

ITS sites have been observed in number of different species but have not been comprehensively characterized within a fully assembled metazoan genome. We located over one thousand ITS sites in the *C. elegans* genome and found a conspicuous depletion of ITS sites at exons and UTRs and enrichment at introns. This localization pattern is not a general feature of hexanulceotide repeats, and is suggestive of specific formation mechanisms and/or selection pressures involved in ITS formation. These data suggest that ITS sites may be toxic to mRNAs, possibly because RNA composed of telomere repeats may form high order structures such as G-quadruplexes (Fay et al., 2017), and may also contribute to intron evolution. Although the origin of ITSs is uncertain, our data suggest that these genome sequences may function to promote gene silencing when in introns and to promote gene expression during development when in enhancers. Alternatively, it is possible that new ITS sites are created in regions of the genome that are silenced or heterochromatic, and that their association with silencing marks is a consequence of their mechanism of biogenesis. Given the abundance of telomeric siRNAs with mismatches that map to ITS tracts and the depletion of ITSs from 3’UTRs, we favor the possibility that ITS tracts may actively promote transcriptional and post-transcriptional gene silencing, a topic that could be addressed by nuclease-mediated genome engineering. Although telomerase generally functions at telomeres, there is evidence to suggest that its reverse transcriptase activity can result in the insertion of telomeric sequence double-strand breaks, although this generally results in the creation of a new telomere (Churikov and Geli, 2017). Therefore, ITSs may represent a contribution to intron and enhancer evolution by the telomerase enzyme.

We found that ITS sites are a shared feature amongst close relatives of *C. elegans*, although their compositions bare some striking differences. ITSs in both *C. briggsae* and *C. remanei* are rarer and generally more degenerate than in *C. elegans*, although both species contain examples of ITS sites with perfect telomere homology. It is intriguing that there should be such a discrepancy in ITS divergence between these species, and this observation suggests differing selective pressures. Although it may be a coincidence that perfect telomeric small RNAs are far more abundant in *C. briggsae* and C. *remanei*, it is also possible that these RNAs are functionally linked to ITS formation and/or evolution. Telomeric small RNAs may actively inhibit formation or expansion of ITS sites, meaning that their very low levels in *C. elegans* could have led to an increasing number of ITSs in the *C. elegans* genome, many of which have been formed relatively recently.

## Materials and Methods

### Analysis of high throughput sequencing data

Publicly available RNA-seq datasets were download from the Gene Expression Omnibus (https://www.ncbi.nlm.nih.gov/geo/) (see supplemental file 3 for library information). Adapter trimming was performed as required using the bbduk.sh script from the bbmap suite3 and custom scripts. Reads were then converted to fasta format and mapped to the *C. elegans* genome (WS251) using Butter (https://www.biorxiv.org/content/10.1101/007427v1). To detect telomere-mapping sRNAs, reads were mapped to seven repeats of perfect telomere sequence (TTAGGC) with Bowtie (Langmead et al., 2009), allowing for up to three mismatches. Sequence pipelines were built using the Snakemake framework (Koster and Rahmann, 2012).

### Genomewide mapping of ITS sites

Bowtie was used to find all instances of the TTAGGC hexamer sequence in the genome. Hexamers located within 100 nucleotides of each other and situated on the same strand were merged into clusters using bedtools (Quinlan and Hall, 2010) and clusters containing four or TTAGGC hexamers were retained. Telomeres or potential telomeres, defined as any ITS that begins or ends within 50 nucleotides of the end of a contig, were removed. In the cases of clashes in which ITSs on opposite strands overlap with one another, the ITS site with the most repeats was retained. Homology scores were assigned to the final set of ITS sites by performing local sequence alignment between each ITS and a stretch of telomere sequence of the same length using the EMBOSS water algorithm http://emboss.sourceforge.net/apps/release/6.6/emboss/apps/water.html.

The method was performed on the *C. elegans* (WS251) *C. briggsae* (WS263) and *C. remanei* (WS263) genomes.

### Analysis of genomic features at ITS sites

Genomic features such as introns, exons etc. were assigned to ITS sites using the GenomicRanges R package [23950696]. Assignment of a particular genomic feature required that at least 50% of the length of the ITS site overlapped with this feature.

Chromatin state data was obtained from http://compbio.med.harvard.edu/modencode/webpage/hihmm/. Chromatin states were assigned to ITS sites using bedtools, requiring that at least 50% of the length of the ITS overlap with the particular chromatin state. To determine enrichment of particular chromatin states in the ITS set, 10000 simulations were performed in which the ITS set was permuted using bedtools shuffle, resulting in a set of random genomic regions with the same lengths as the real ITS set. The regions were divided into intergenic and intronic regions and were assigned chromatin states. Enrichment for a particular chromatin state was calculated by dividing the number of ITS sites with that chromatin state by the average number of sites with the same state in the simulated data. P-values were obtained by determining the percentage of simulations in which the number of sites that were assigned a particular chromatin state was equal to or more (for enrichment) or equal to or fewer (for depletion) than the real ITS set. P-values were adjusted using Benjamini–Hochberg procedure (FDR). Given that 10000 simulations were performed, the rarest possible event that can be detected is one that occurs one in 10000 simulations, corresponding to a rate of 0.0001. After Benjamini–Hochberg correction, this value becomes 0.00048. Therefore, “p < 0.00048” was assigned in the cases in which the number of ITS sites with a particular chromatin state was more extreme than any of the 10000 simulations.

Lists of orthologous genes between *C. elegans, C. briggsae* and *C. remanei* were obtained from Parasite.

### Statistics and data visualization

Analysis of sequencing data was performed using the R statistical computing environment. Genome regions were visualized using the Gviz R package (Hahne and Ivanek, 2016). GO term enrichment analysis was performed using the RDAVIDWebService R package (Fresno and Fernandez, 2013).

### Southern Blotting

Genomic DNA was prepared from *C. elegans* strain Bristol N2, *C. remanei* strains PB4641, EM464 and SB146, and *C. briggsae* strains MK104 and AF16, which were grown at 20°C on NGM plates. Southern blotting was performed using genomic DNA prepared from a Gentra kit and a digoxygenin-labelled TTAGGC probe, as previously described (Meier et al., 2009).

## Acknowledgements

We thank Yun Li, Will Valdar and Erin Osborne for advice concerning genome analysis. This work was supported by NIH R01 GM066228 to SA.

## Author Contributions

S.F. and S.A. designed the experiments. S.F. and E.H.L.-S. carried out the experiments, S.F. performed computational analysis and E.H.L.-S. performed Southern blotting, S.F. and S.A. wrote the manuscript.

---

**Table S1.** Chromatin state enrichment for ITS sites based on simulations

| stage | state | feature | count | enrichment | p-value |
| --- | --- | --- | --- | --- | --- |
| ee | Enhancer | intergenic | 59 | 0.6778360552 | 0.00048 |
| ee | Enhancer | intron | 55 | -0.1570874597 | 0.3279272727 |
| ee | Gene body | intergenic | 57 | -0.4570613948 | 0.0006 |
| ee | Gene body | intron | 209 | -0.3705784898 | 0.0066857143 |
| ee | Heterochromatin | intergenic | 34 | -0.0892991405 | 0.3279272727 |
| ee | Heterochromatin | intron | 119 | 1.2496763643 | 0.0075 |
| ee | Low signal | intergenic | 182 | 0.0156254583 | 0.4227130435 |
| ee | Low signal | intron | 227 | -0.449121839 | 0.0075 |
| ee | Polycomb | intergenic | 52 | 1.1496090691 | 0.0003 |
| ee | Polycomb | intron | 160 | 2.1278433535 | 0.0003 |
| ee | Promoter | intergenic | 2 | -3.1452021959 | 0.0003 |
| ee | Promoter | intron | 9 | -1.2488723695 | 0.0219789474 |
| l3 | Enhancer | intergenic | 98 | 0.9958072399 | 0.0003 |
| l3 | Enhancer | intron | 66 | -0.4703345716 | 0.19668 |
| l3 | Gene body | intergenic | 22 | -2.0003681619 | 0.0003 |
| l3 | Gene body | intron | 214 | -0.4505126037 | 0.0051692308 |
| l3 | Heterochromatin | intergenic | 46 | -0.0289542175 | 0.4245 |
| l3 | Heterochromatin | intron | 172 | 1.1047287454 | 0.0168 |
| l3 | Low signal | intergenic | 102 | -0.5127075469 | 0.0006 |
| l3 | Low signal | intron | 97 | -1.0541539975 | 0.00048 |
| l3 | Polycomb | intergenic | 116 | 2.0772017409 | 0.0003 |
| l3 | Polycomb | intron | 228 | 2.1816948244 | 0.0003 |
| l3 | Promoter | intergenic | 2 | -2.4362512894 | 0.0003 |
| I3 | Promoter | intron | 2 | -2.4225137827 | 0.0196 |

**Table S2.**
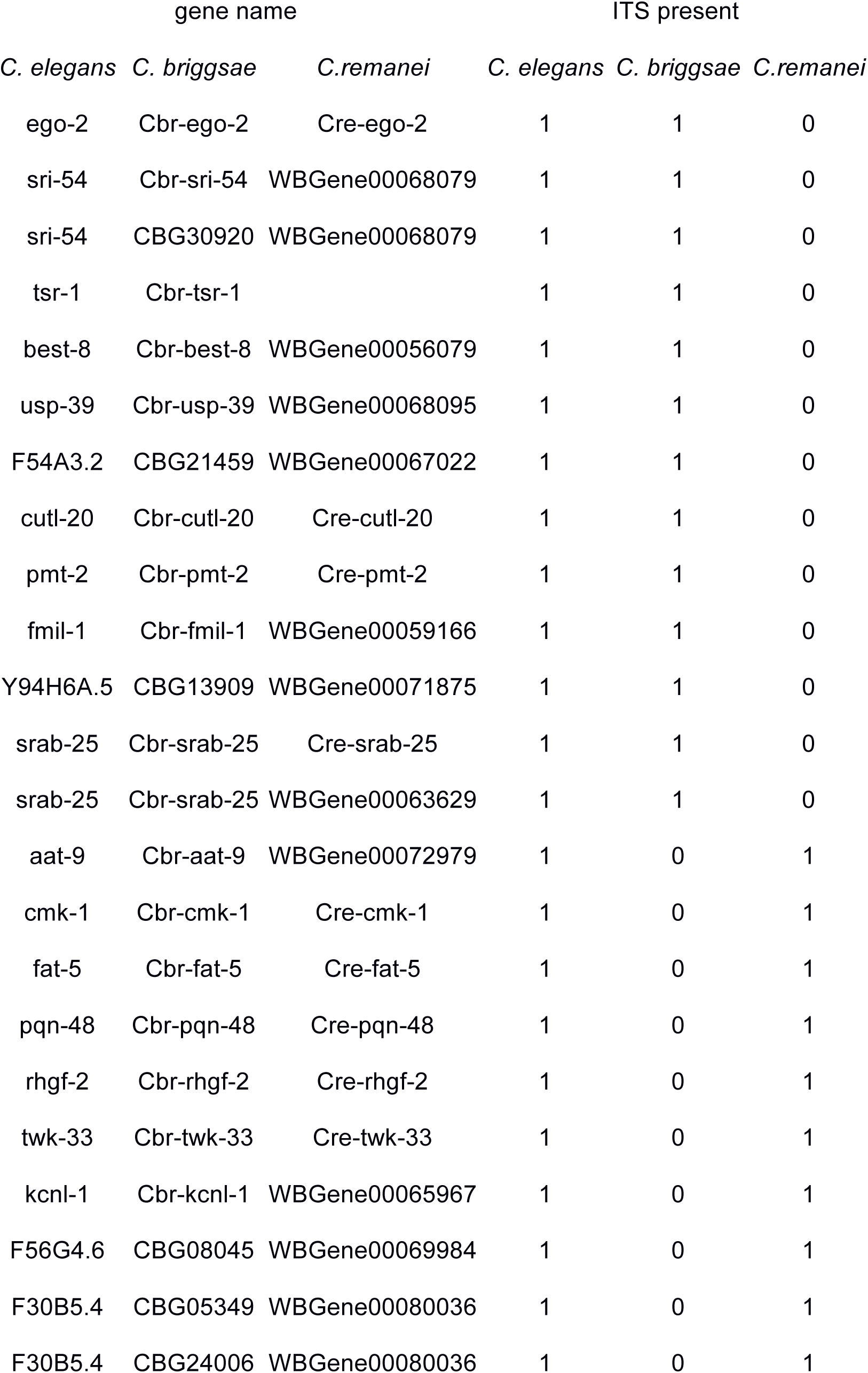
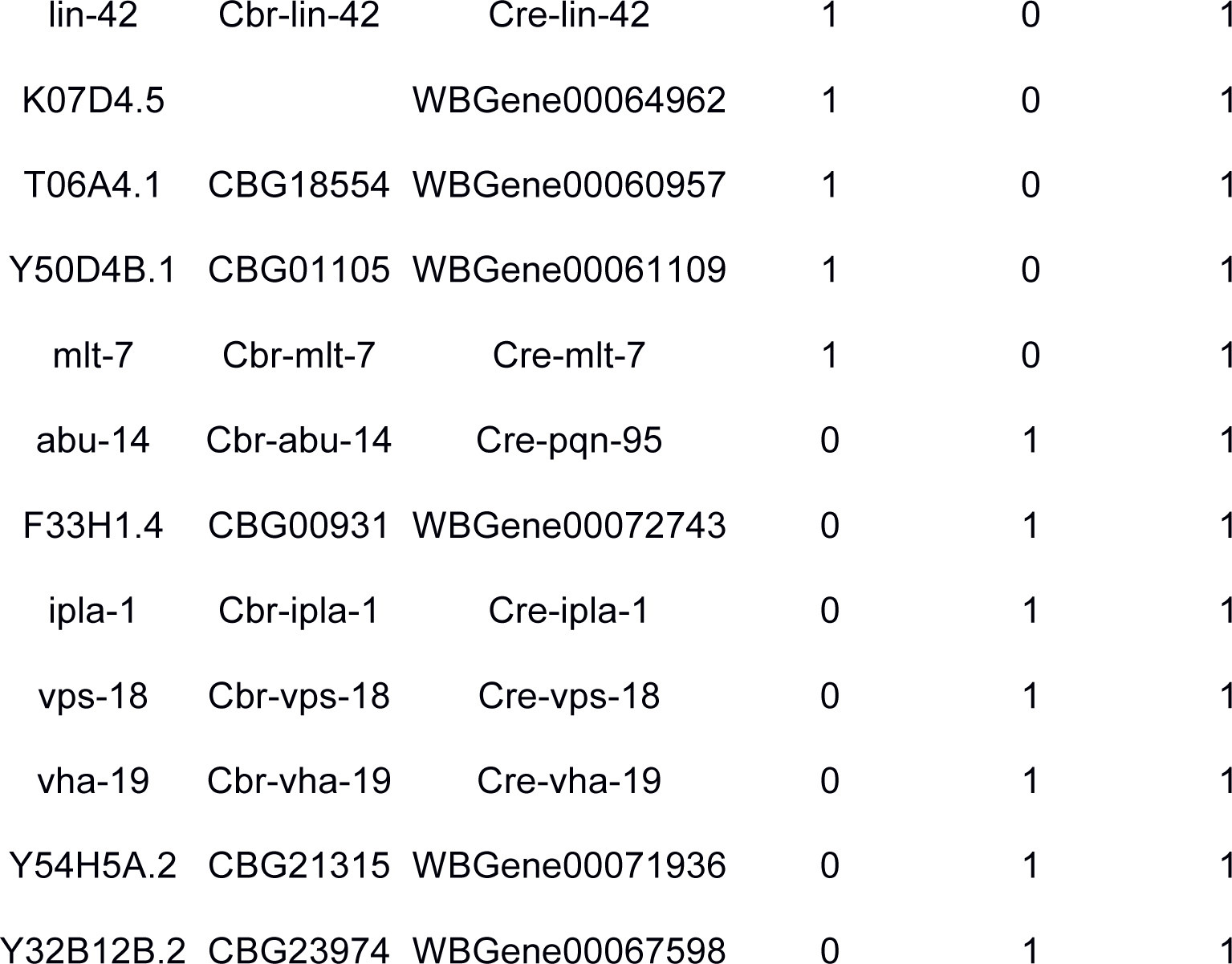
Genes which contain ITS sites in more than one species

**Table S3.**
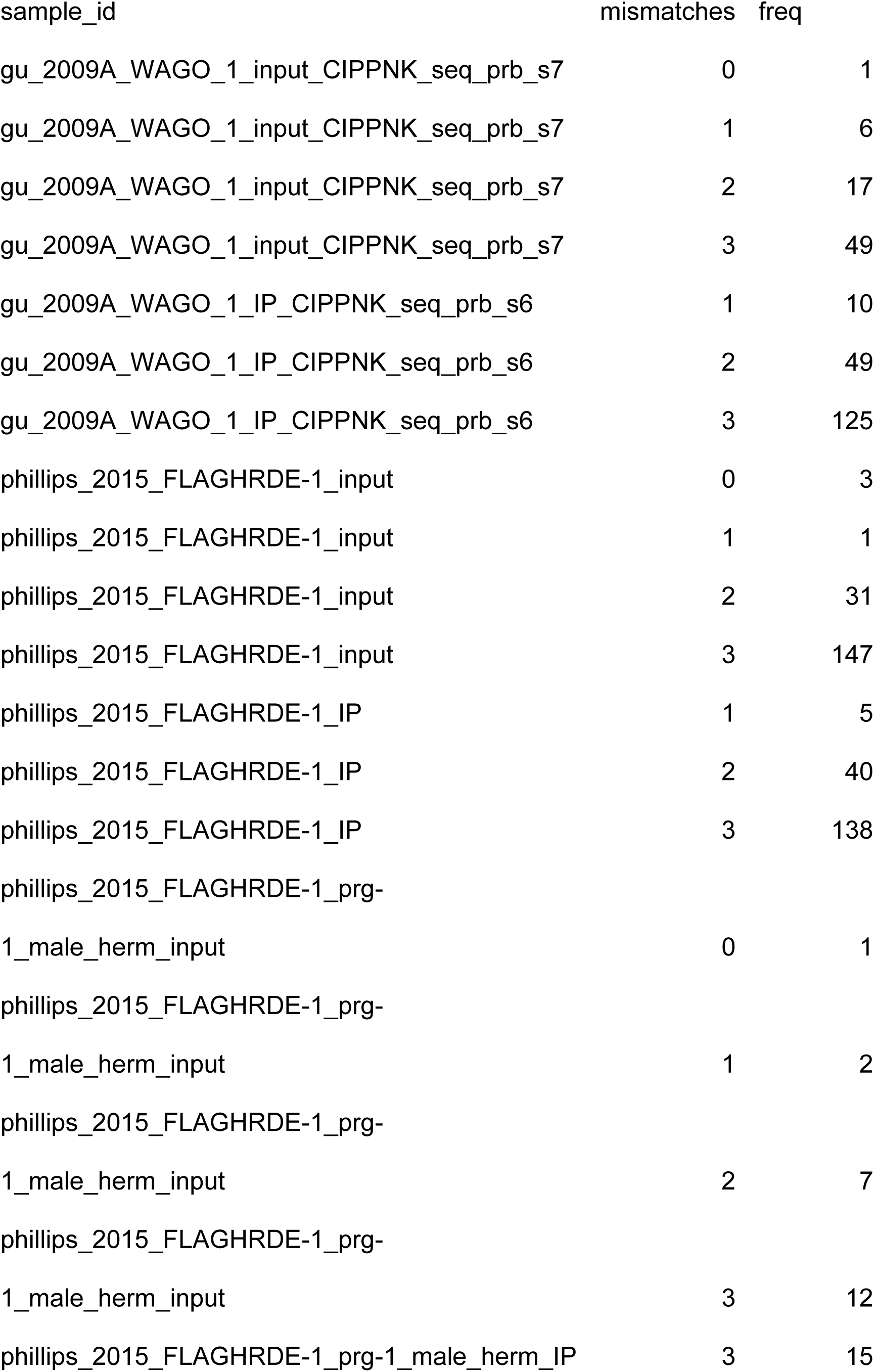
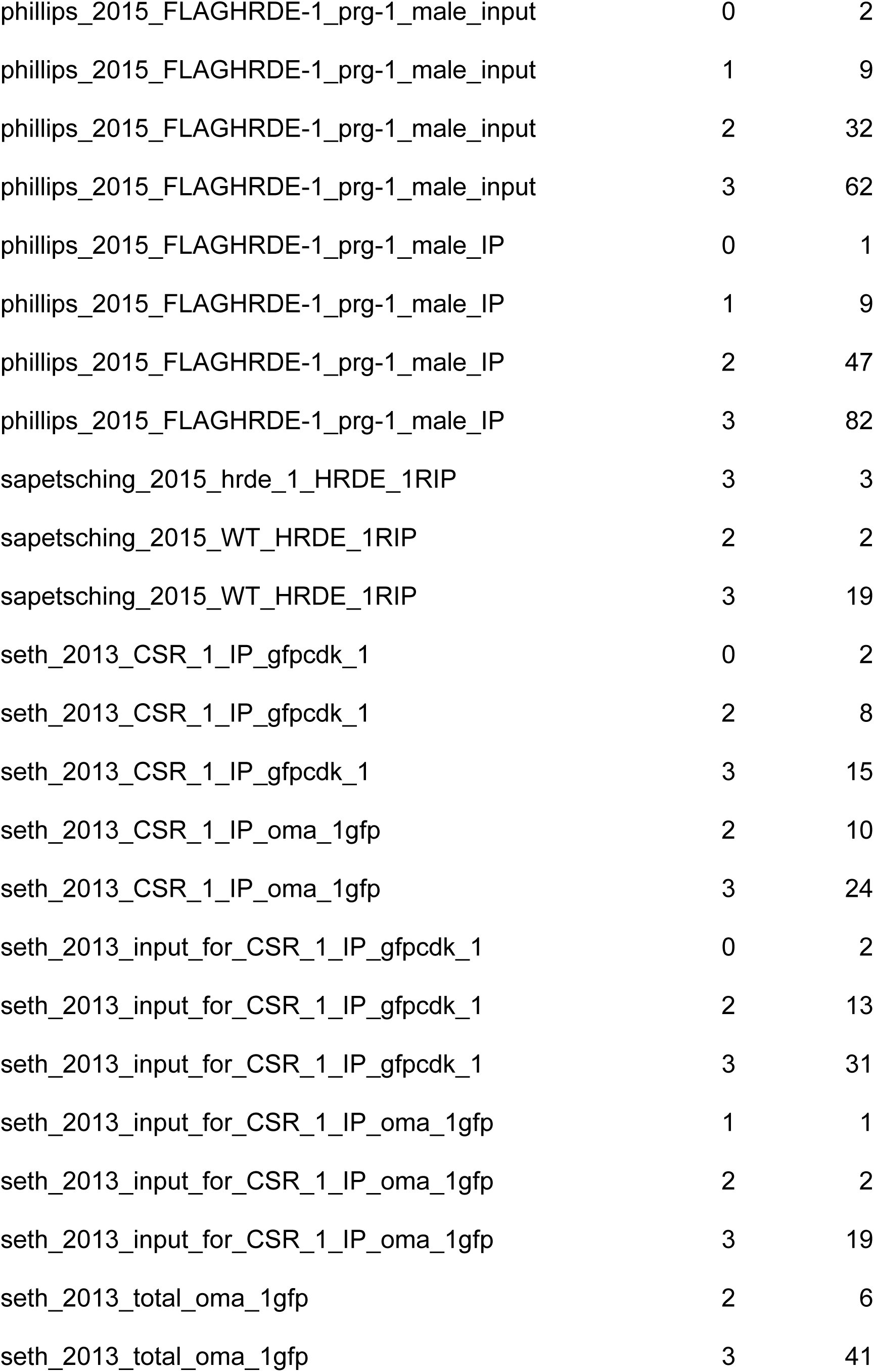
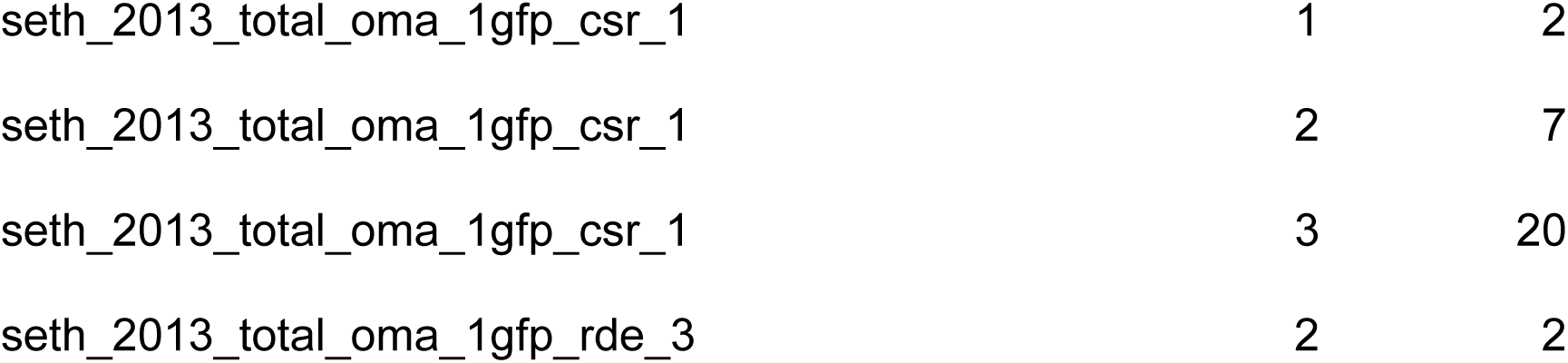
Total telomere sRNAs from input and IP datasets.

**Table S4.**
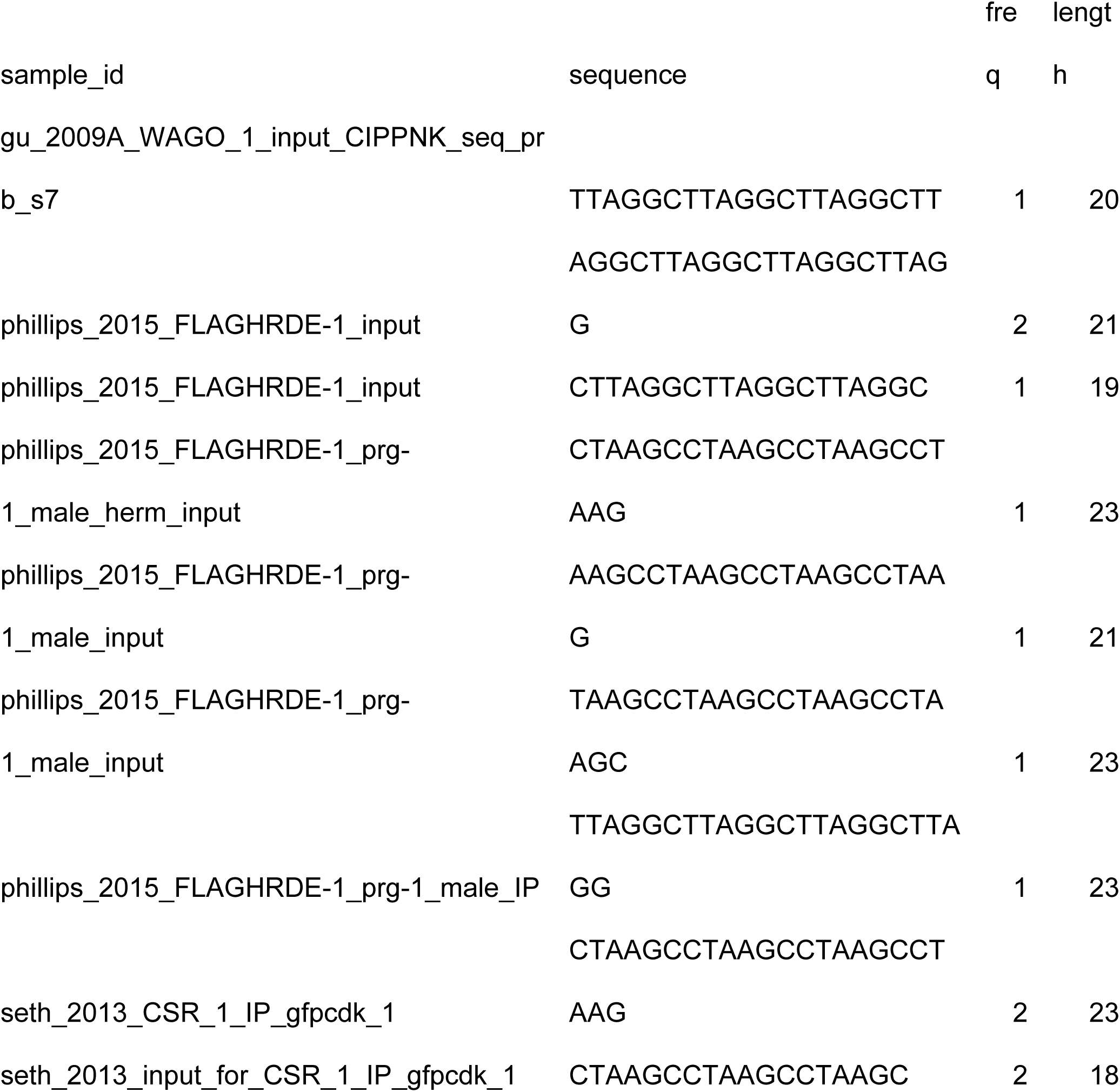
Perfect telomeric sRNAs from input and IP datasets.

**Table S5.** Telomeric reads from 77 small RNA libraries from mouse with 0, 1, 2 or 3 mismatches.

| sample_id | 0 | 1 | 2 | 3 |
| --- | --- | --- | --- | --- |
| miwi2HET_E165_Testis_smallRNA | 36 | 7 | 10 | 59 |
| miwi2HET_P10_Testis_smallRNA | 2 | 1 | 2 | 37 |
| miwi2KO_E165_Testis_smallRNA | 91 | 13 | 11 | 70 |
| BmTdrd12_IP | 0 | 0 | 0 | 3 |
| HA_BmTdrd12_IP | 0 | 0 | 0 | 7 |
| MILI_IP | 0 | 1 | 2 | 9 |
| MILI_IP_ | 0 | 2 | 3 | 6 |
| TDRD12_IP_ | 0 | 4 | 1 | 9 |
| Total_small_RNA_14_40_nt_rep1 | 0 | 0 | 1 | 37 |
| Total_small_RNA_14_40_nt_rep2 | 0 | 0 | 1 | 43 |
| trimmed_10_dpp | 1 | 0 | 1 | 0 |
| trimmed_15_dpp | 2 | 2 | 4 | 3 |
| trimmed_18_dpp | 0 | 0 | 1 | 1 |
| trimmed_40_dpp | 2 | 3 | 3 | 9 |
| trimmed_6_dpp | 2 | 0 | 0 | 1 |
| trimmed_Input_Miwi_ADH__1 | 1 | 0 | 0 | 0 |
| trimmed_LZ | 1 | 0 | 0 | 0 |
| trimmed_MIWI_IP_Miwi__1 | 1 | 0 | 0 | 0 |
| trimmed_MIWI_IP_Miwi__2 | 2 | 0 | 0 | 1 |
| trimmed_MIWI_IP_Miwi___2 | 0 | 0 | 0 | 1 |
| trimmed_MIWI_IP_Miwi_ADH__1 | 0 | 0 | 1 | 0 |
| trimmed_Miwi__late_sc_rep1 | 0 | 0 | 0 | 1 |
| MVH__adultMILI_IPlonger_RNAs | 0 | 0 | 0 | 1 |
| MVH__adultMILI_IPpiRNAs | 0 | 0 | 0 | 1 |
| MVH__adultMIWI_IPlonger_RNAs | 0 | 0 | 0 | 10 |
| MVH__adultMIWI_IPpiRNAs | 0 | 0 | 0 | 23 |
| MVH_adultMVH_IPlonger_RNAs | 2 | 0 | 1 | 20 |
| MVH_adultMVH_IPpiRNAs | 0 | 1 | 0 | 15 |
| MVH_KladultMILI_IPlonger_RNAs | 0 | 0 | 0 | 1 |
| MVH_KladultMILI_IPpiRNAs | 0 | 0 | 0 | 2 |
| MVH_KladultMVH_IPlonger_RNAsrep1 | 0 | 0 | 0 | 3 |
| MVH_KladultMVH_IPlonger_RNAsrep3 | 0 | 0 | 1 | 3 |
| MVH_KladultMVH_IPpiRNAsrep1 | 1 | 0 | 0 | 3 |
| MVH_KladultMVH_IPpiRNAsrep2 | 0 | 0 | 0 | 1 |
| MVH__KIP0MILI_IPlonger_RNAs | 0 | 0 | 1 | 6 |
| MVH_KIP0MILI_IPlonger_RNAs | 0 | 0 | 1 | 5 |
| MVH__KIP0MILI_IPpiRNAs | 2 | 1 | 3 | 14 |
| MVH_KIP0MILI_IPpiRNAs | 0 | 2 | 3 | 25 |
| MVH_KIP0MIWI2_IPlonger_RNAs | 0 | 0 | 1 | 32 |
| MVH_KIP0MIWI2_IPpiRNAs | 0 | 1 | 3 | 129 |
| MVH__P0MILI_IPpiRNAs | 0 | 1 | 0 | 12 |
| MVH__P0MIWI2_IPlonger_RNAs | 0 | 0 | 2 | 16 |
| MVH__P0MIWI2_IPpiRNAs | 0 | 2 | 0 | 32 |
| Rosa26_pi_reporterMVH__KIP0MILI_IPlonger_RNAs | 0 | 0 | 0 | 5 |
| Rosa26_pi_reporterMVH_KIP0MILI_IPlonger_RNAs | 1 | 0 | 0 | 4 |
| Rosa26_pi_reporterMVH__KIP0MILI_IPpiRNAs | 2 | 5 | 3 | 32 |
| Rosa26_pi_reporterMVH_KIP0MILI_IPpiRNAs | 2 | 3 | 4 | 40 |
| Rosa26_pi_reporterMVH__P0MILI_IPlonger_RNA<br>s | 1 | 1 | 5 | 8 |
| Rosa26_pi_reporterMVH__P0MILI_IPlonger_RNAs | 0 | 0 | 1 | 10 |
| Rosa26_pi_reporterMVH__P0MILI_IPpiRNAs | 1 | 2 | 5 | 53 |
| Rosa26_pi_reporterMVH_P0MILI_IPpiRNAs | 3 | 3 | 7 | 22 |
| Tdrd9_KIKIP0MILI_IPpiRNAs | 0 | 1 | 0 | 3 |
| Tdrd9_KIKIP0MIWI2_IPpiRNAs | 0 | 0 | 1 | 14 |
| Tdrd9_KIP0MILI_IPpiRNAs | 0 | 0 | 2 | 5 |
| Tdrd9_KIP0MIWI2_IPpiRNAs | 0 | 0 | 1 | 6 |
| Tdrd9__P0MILI_IPpiRNAs | 0 | 0 | 0 | 1 |
| Tdrd9__P0MIWI2_IPpiRNAs | 0 | 0 | 2 | 6 |

**Table S6.**
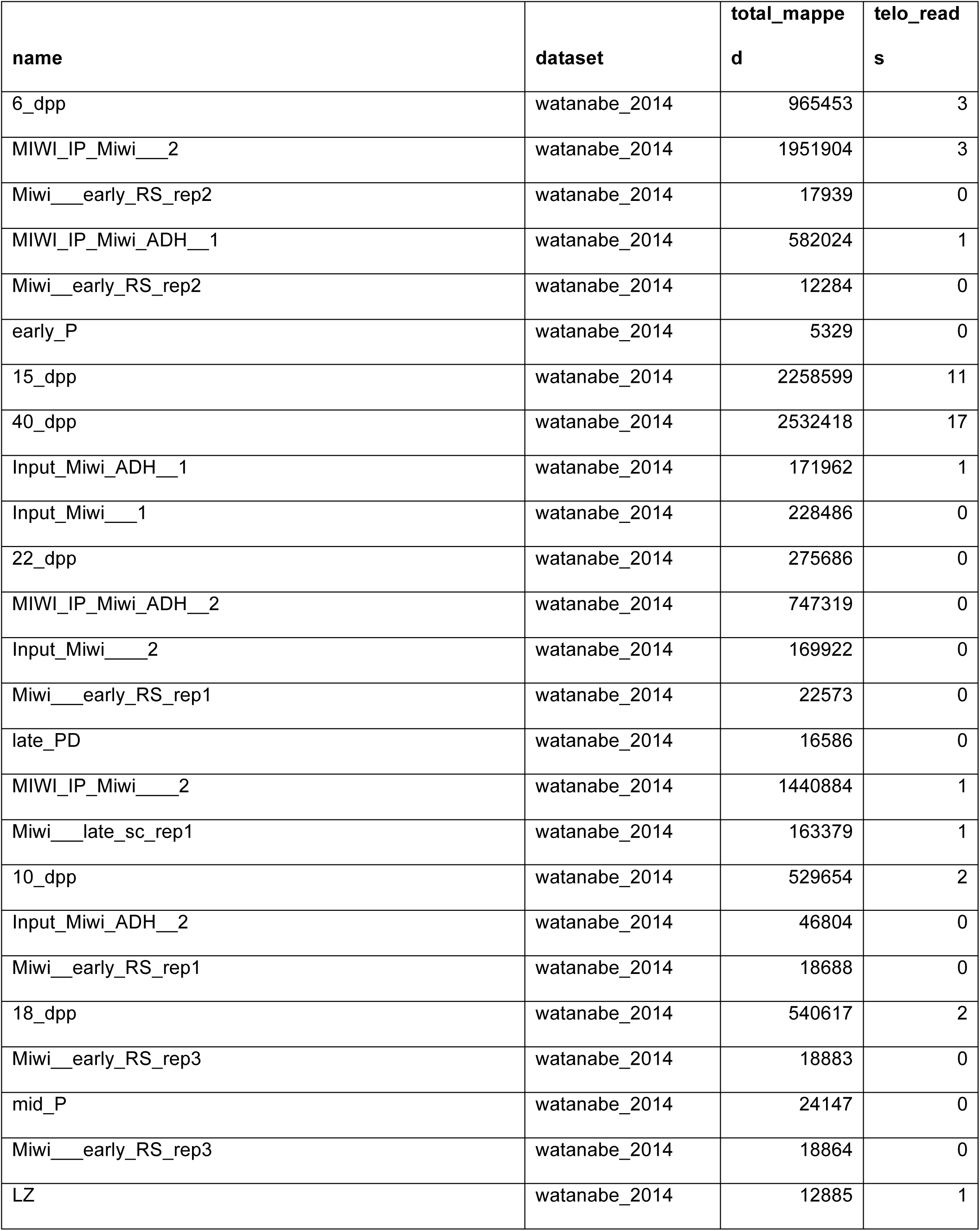
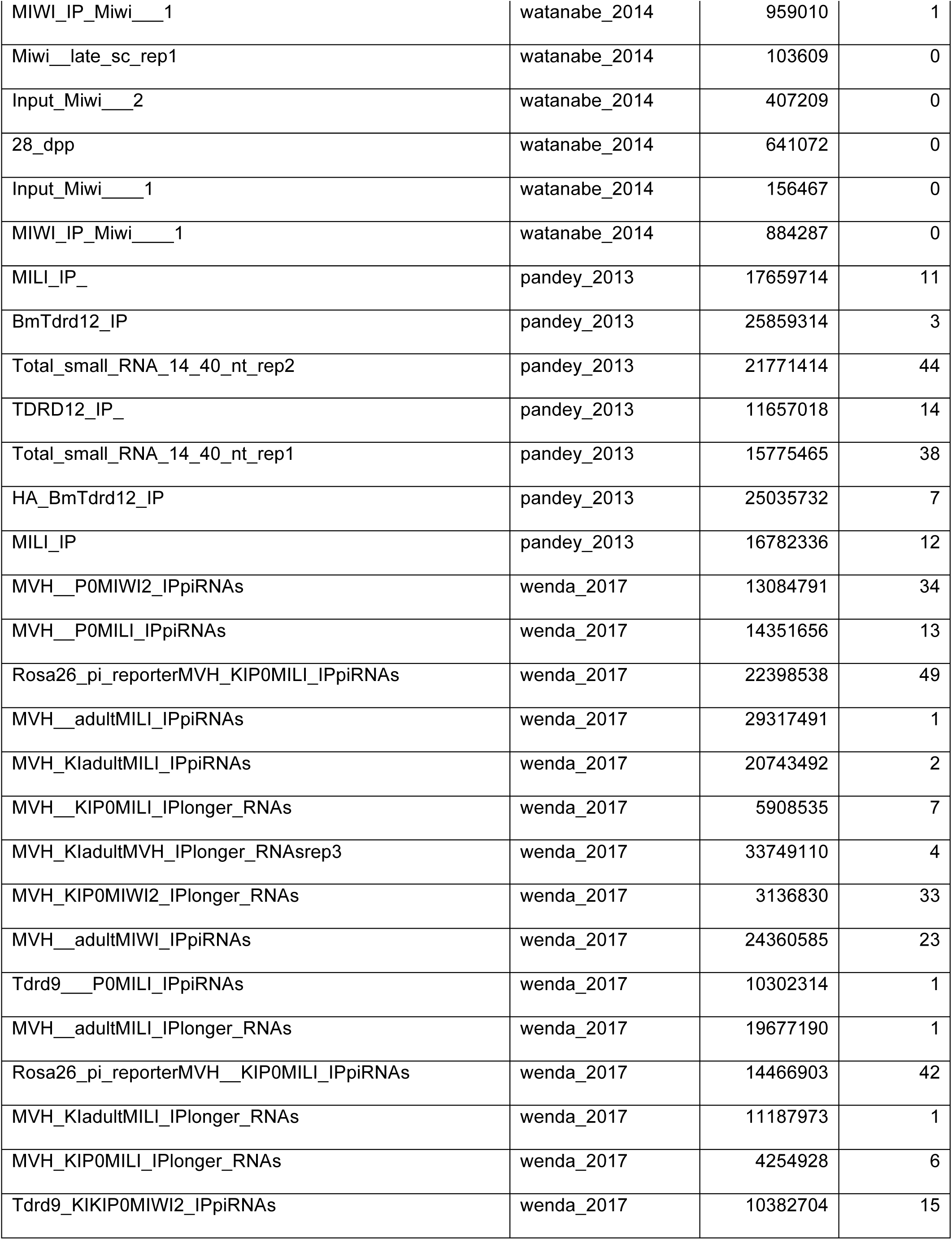
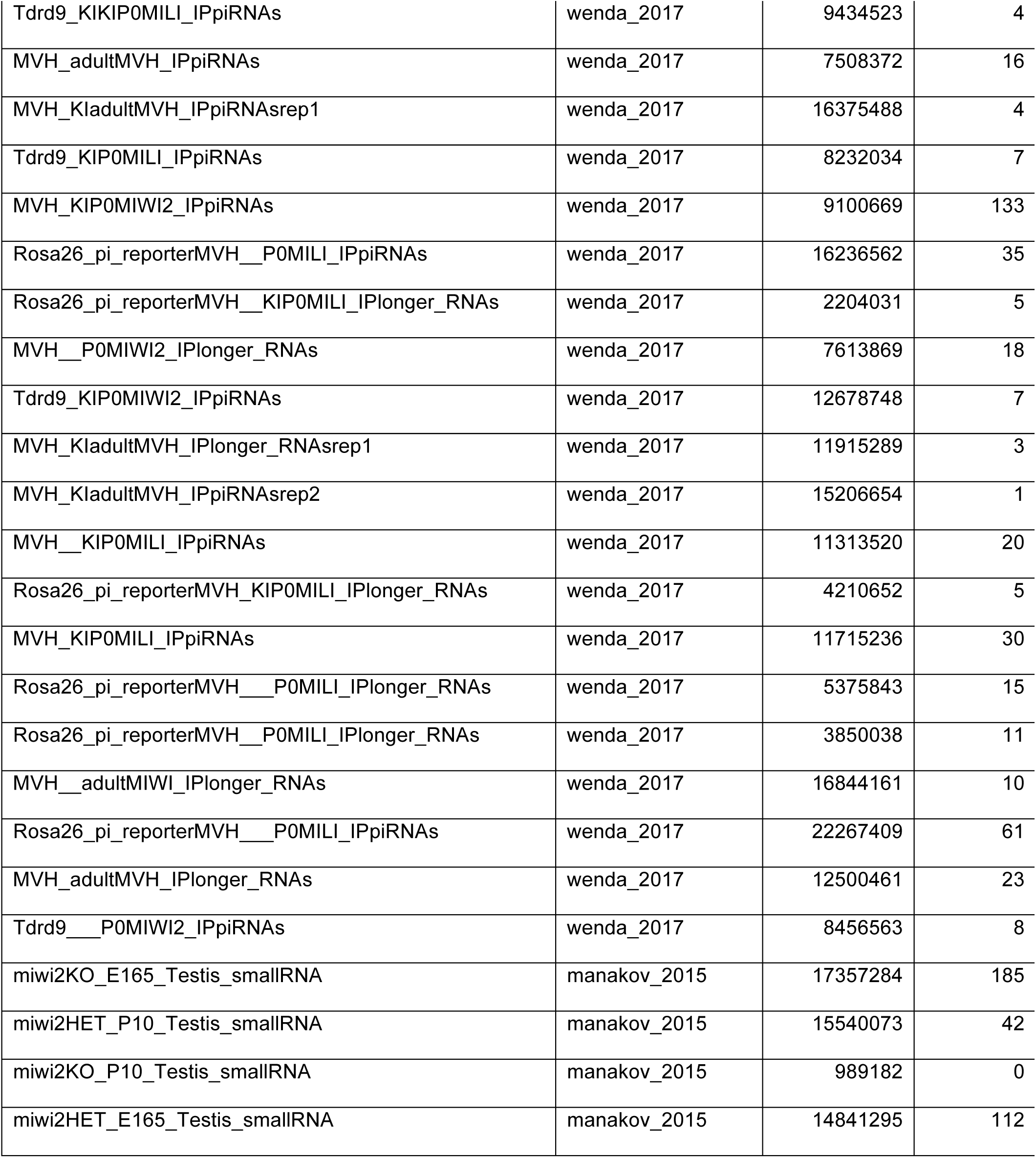
Sample information for mouse small RNA analysis.

| **name** | **dataset** | **total_mappe**<br>**d** | **telo_read**<br>**s** |
| --- | --- | --- | --- |
| 6_dpp | watanabe_2014 | 965453 | 3 |
| MIWI_IP_Miwi___2 | watanabe_2014 | 1951904 | 3 |
| Miwi___early_RS_rep2 | watanabe_2014 | 17939 | 0 |
| MIWI_IP_Miwi_ADH___1 | watanabe_2014 | 582024 | 1 |
| Miwi___early_RS_rep2 | watanabe_2014 | 12284 | 0 |
| early_P | watanabe_2014 | 5329 | 0 |
| 15_dpp | watanabe_2014 | 2258599 | 11 |
| 40_dpp | watanabe_2014 | 2532418 | 17 |
| Input_Miwi_ADH___1 | watanabe_2014 | 171962 | 1 |
| Input_Miwi___1 | watanabe_2014 | 228486 | 0 |
| 22_dpp | watanabe_2014 | 275686 | 0 |
| MIWI_IP_Miwi_ADH___2 | watanabe_2014 | 747319 | 0 |
| Input_Miwi___2 | watanabe_2014 | 169922 | 0 |
| Miwi___early_RS_rep1 | watanabe_2014 | 22573 | 0 |
| late_PD | watanabe_2014 | 16586 | 0 |
| MIWI_IP_Miwi___2 | watanabe_2014 | 1440884 | 1 |
| Miwi___late_sc_rep1 | watanabe_2014 | 163379 | 1 |
| 10_dpp | watanabe_2014 | 529654 | 2 |
| Input_Miwi_ADH___2 | watanabe_2014 | 46804 | 0 |
| Miwi___early_RS_rep1 | watanabe_2014 | 18688 | 0 |
| 18_dpp | watanabe_2014 | 540617 | 2 |
| Miwi___early_RS_rep3 | watanabe_2014 | 18883 | 0 |
| mid_P | watanabe_2014 | 24147 | 0 |
| Miwi___early_RS_rep3 | watanabe_2014 | 18864 | 0 |
| LZ | watanabe_2014 | 12885 | 1 |
| MIWI_IP_Miwi___1 | watanabe_2014 | 959010 | 1 |
| Miwi__late_sc_rep1 | watanabe_2014 | 103609 | 0 |
| Input_Miwi___2 | watanabe_2014 | 407209 | 0 |
| 28_dpp | watanabe_2014 | 641072 | 0 |
| Input_Miwi___1 | watanabe_2014 | 156467 | 0 |
| MIWI_IP_Miwi___1 | watanabe_2014 | 884287 | 0 |
| MILI_IP_ | pandey_2013 | 17659714 | 11 |
| BmTdrd12_IP | pandey_2013 | 25859314 | 3 |
| Total_small_RNA_14_40_nt_rep2 | pandey_2013 | 21771414 | 44 |
| TDRD12_IP_ | pandey_2013 | 11657018 | 14 |
| Total_small_RNA_14_40_nt_rep1 | pandey_2013 | 15775465 | 38 |
| HA_BmTdrd12_IP | pandey_2013 | 25035732 | 7 |
| MILI_IP | pandey_2013 | 16782336 | 12 |
| MVH__P0MIWI2_IPpiRNAs | wenda_2017 | 13084791 | 34 |
| MVH__P0MILI_IPpiRNAs | wenda_2017 | 14351656 | 13 |
| Rosa26_pi_reporterMVH_KIP0MILI_IPpiRNAs | wenda_2017 | 22398538 | 49 |
| MVH__adultMILI_IPpiRNAs | wenda_2017 | 29317491 | 1 |
| MVH_KladultMILI_IPpiRNAs | wenda_2017 | 20743492 | 2 |
| MVH__KIP0MILI_IPlonger_RNAs | wenda_2017 | 5908535 | 7 |
| MVH_KladultMVH_IPlonger_RNAsrep3 | wenda_2017 | 33749110 | 4 |
| MVH_KIP0MIWI2_IPlonger_RNAs | wenda_2017 | 3136830 | 33 |
| MVH__adultMIWI_IPpiRNAs | wenda_2017 | 24360585 | 23 |
| Tdrd9__P0MILI_IPpiRNAs | wenda_2017 | 10302314 | 1 |
| MVH__adultMILI_IPlonger_RNAs | wenda_2017 | 19677190 | 1 |
| Rosa26_pi_reporterMVH__KIP0MILI_IPpiRNAs | wenda_2017 | 14466903 | 42 |
| MVH_KladultMILI_IPlonger_RNAs | wenda_2017 | 11187973 | 1 |
| MVH_KIP0MILI_IPlonger_RNAs | wenda_2017 | 4254928 | 6 |
| Tdrd9_KIKIP0MIWI2_IPpiRNAs | wenda_2017 | 10382704 | 15 |
| Tdrd9_KIKIP0MILI_IPpiRNAs | wenda_2017 | 9434523 | 4 |
| MVH_adultMVH_IPpiRNAs | wenda_2017 | 7508372 | 16 |
| MVH_KladultMVH_IPpiRNAsrep1 | wenda_2017 | 16375488 | 4 |
| Tdrd9_KIP0MILI_IPpiRNAs | wenda_2017 | 8232034 | 7 |
| MVH_KIP0MIWI2_IPpiRNAs | wenda_2017 | 9100669 | 133 |
| Rosa26_pi_reporterMVH__P0MILI_IPpiRNAs | wenda_2017 | 16236562 | 35 |
| Rosa26_pi_reporterMVH__KIP0MILI_IPlonger_RNAs | wenda_2017 | 2204031 | 5 |
| MVH__P0MIWI2_IPlonger_RNAs | wenda_2017 | 7613869 | 18 |
| Tdrd9_KIP0MIWI2_IPpiRNAs | wenda_2017 | 12678748 | 7 |
| MVH_KladultMVH_IPlonger_RNAsrep1 | wenda_2017 | 11915289 | 3 |
| MVH_KladultMVH_IPpiRNAsrep2 | wenda_2017 | 15206654 | 1 |
| MVH__KIP0MILI_IPpiRNAs | wenda_2017 | 11313520 | 20 |
| Rosa26_pi_reporterMVH_KIP0MILI_IPlonger_RNAs | wenda_2017 | 4210652 | 5 |
| MVH_KIP0MILI_IPpiRNAs | wenda_2017 | 11715236 | 30 |
| Rosa26_pi_reporterMVH__P0MILI_IPlonger_RNAs | wenda_2017 | 5375843 | 15 |
| Rosa26_pi_reporterMVH__P0MILI_IPlonger_RNAs | wenda_2017 | 3850038 | 11 |
| MVH__adultMIWI_IPlonger_RNAs | wenda_2017 | 16844161 | 10 |
| Rosa26_pi_reporterMVH__P0MILI_IPpiRNAs | wenda_2017 | 22267409 | 61 |
| MVH_adultMVH_IPlonger_RNAs | wenda_2017 | 12500461 | 23 |
| Tdrd9__P0MIWI2_IPpiRNAs | wenda_2017 | 8456563 | 8 |
| miwi2KO_E165_Testis_smallRNA | manakov_2015 | 17357284 | 185 |
| miwi2HET_P10_Testis_smallRNA | manakov_2015 | 15540073 | 42 |
| miwi2KO_P10_Testis_smallRNA | manakov_2015 | 989182 | 0 |
| miwi2HET_E165_Testis_smallRNA | manakov_2015 | 14841295 | 112 |

**Table S7.**
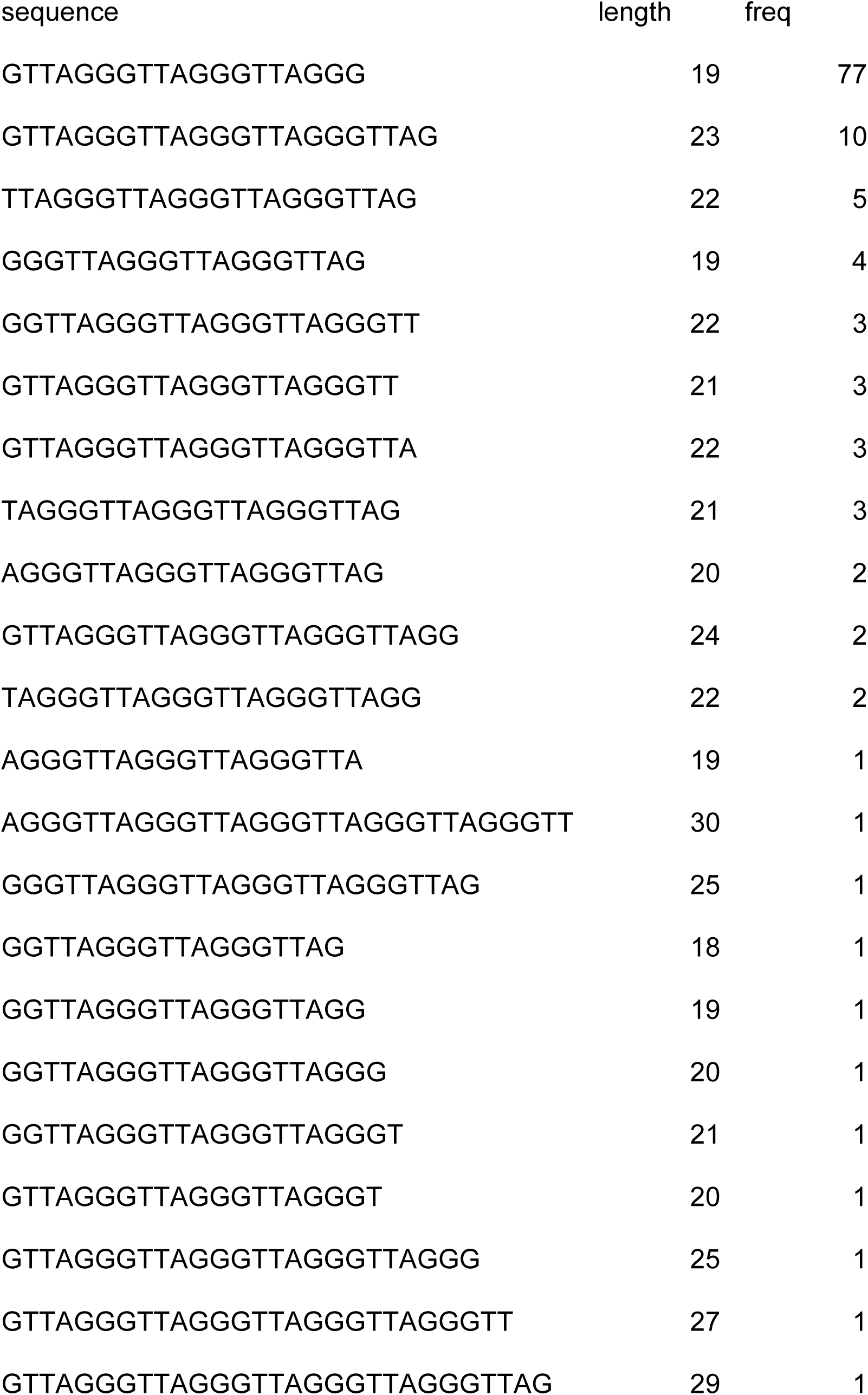
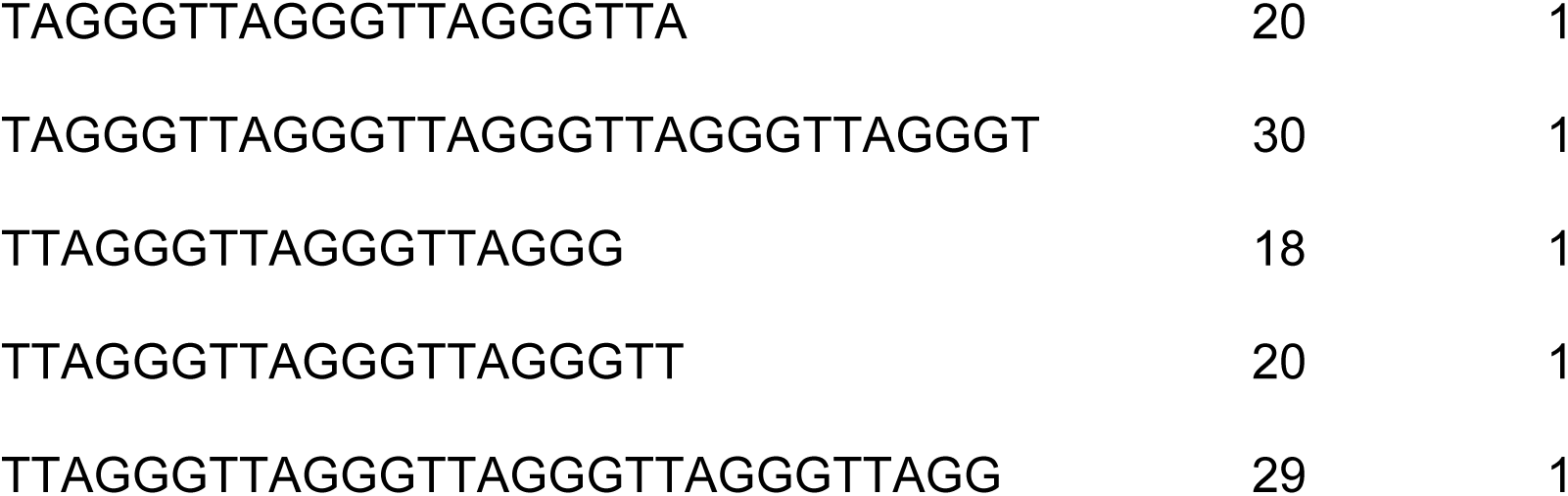
Perfect G-rich telomeric sRNAs from mouse small RNA datasets.

**Table S8.**
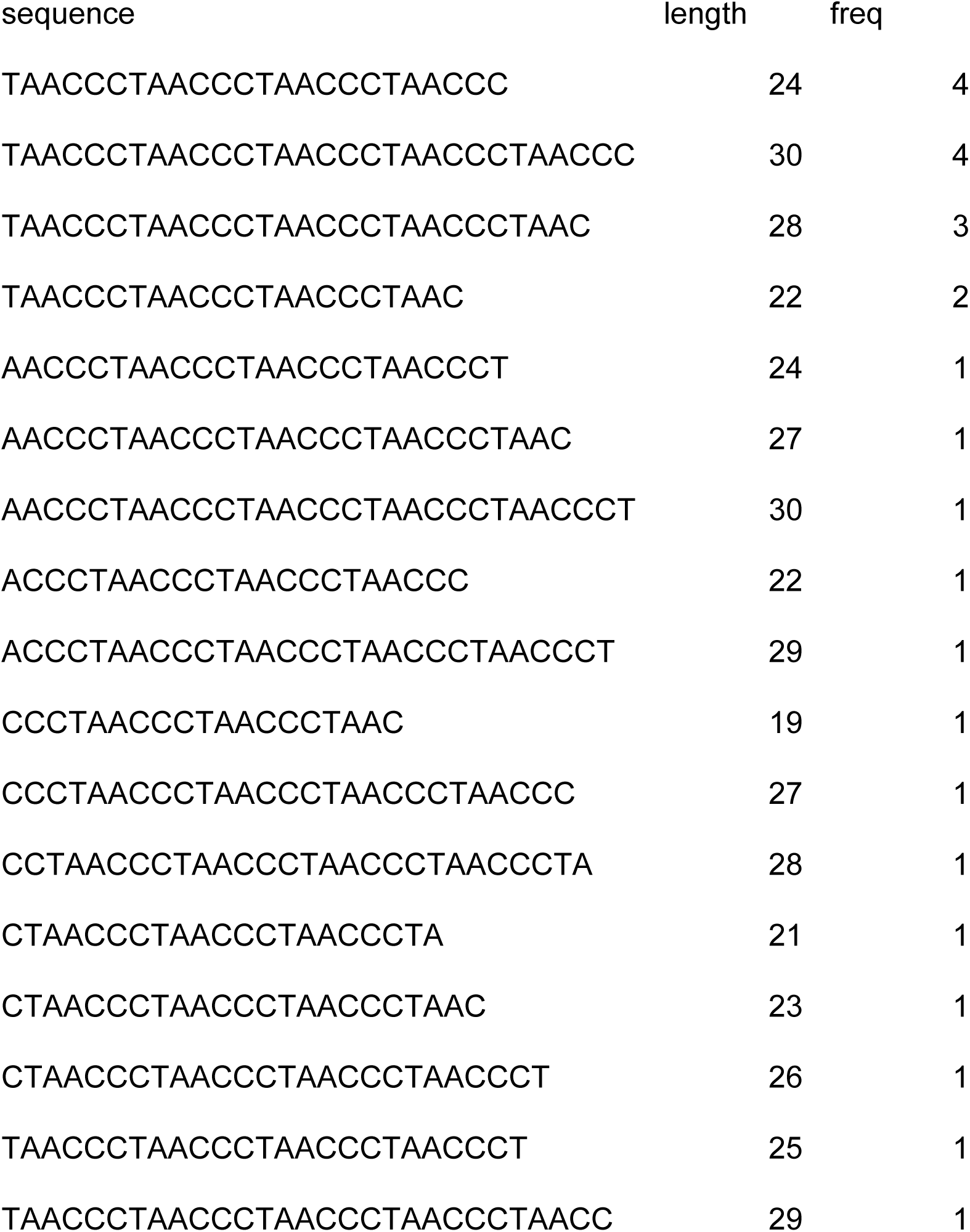
Perfect C-rich telomeric sRNAs from mouse small RNA datasets.

**Table S9.**
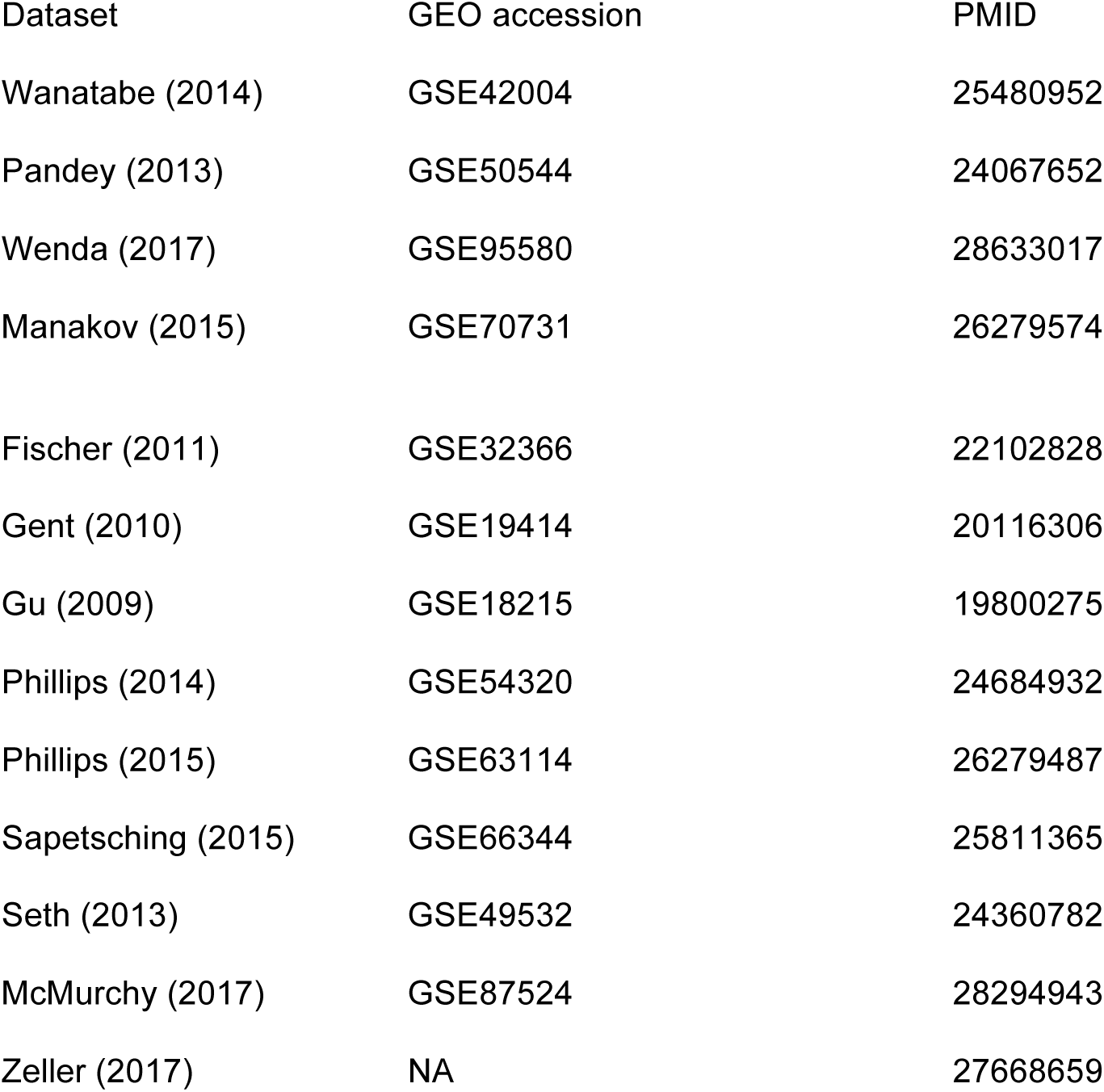
Accession numbers for data sets analyzed in this study. Top 4 for mouse sRNAs and bottom 9 for *C. elegans* sRNAs.

**Figure S1:**
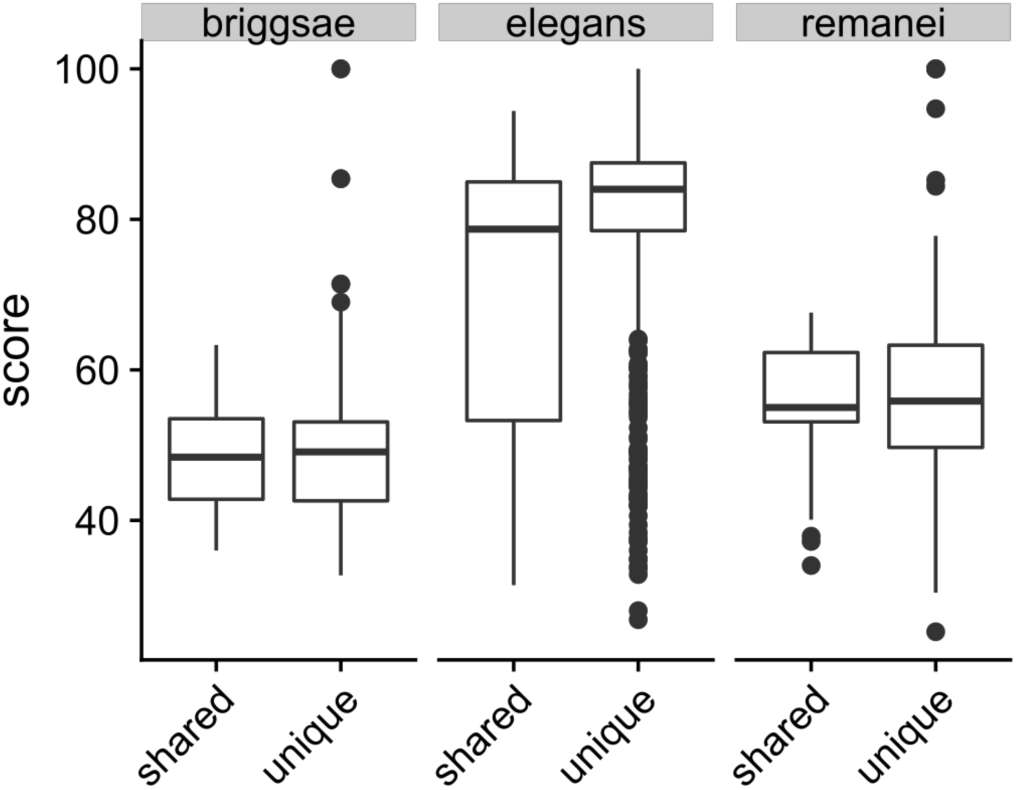
Telomere homology scores for intronic ITS sites broke down by whether or not a homologue of the gene containing the ITS site also contains an ITS site in another species (“shared” versus “unique” respectively).

